# Ocular immune privilege in action: the living eye imposes unique regulatory and anergic gene signatures on uveitogenic T cells

**DOI:** 10.1101/2025.03.01.640701

**Authors:** Zixuan Peng, Vijayaraj Nagarajan, Reiko Horai, Yingyos Jittayasothorn, Mary J. Mattapallil, Rachel R. Caspi

**Affiliations:** Laboratory of Immunology, National Eye Institute, National Institutes of Health, Bethesda, MD, USA; Department of Ophthalmology, Xiangya Hospital, Central South University, Changsha, China

**Keywords:** Ocular immune privilege, autoimmune uveitis, Tregs, T cell anergy, single-cell transcriptomics, immune tolerance

## Abstract

Despite ocular immune privilege, circulating retina-specific T cells can trigger autoimmune uveitis, yet intraocular bleeding—a relatively common event—rarely leads to disease. Using an in vivo immune privilege model, we previously reported that all naïve retina-specific T cells entering the eye become primed *in situ*; about Ob% become FoxpO+ T-regulatory cells (Tregs), while the rest fail to induce pathology. Here, single-cell transcriptomics and functional validation revealed distinct phenotypes in both populations: ocular Tregs were highly suppressive, whereas non-Tregs expressed suppression- and anergy-associated genes and lacked regulatory function. Trajectory analyses suggested that Tregs and anergic cells arise from a common proliferative precursor in parallel, rather than sequentially. Our data indicate a key checkpoint governing the divergence of anergic and regulatory fates. These findings provide molecular-level insights into ocular immune privilege and may inform strategies to silence autoimmune effector cells or reverse T cell unresponsiveness in cancer, vaccination, or chronic infection.

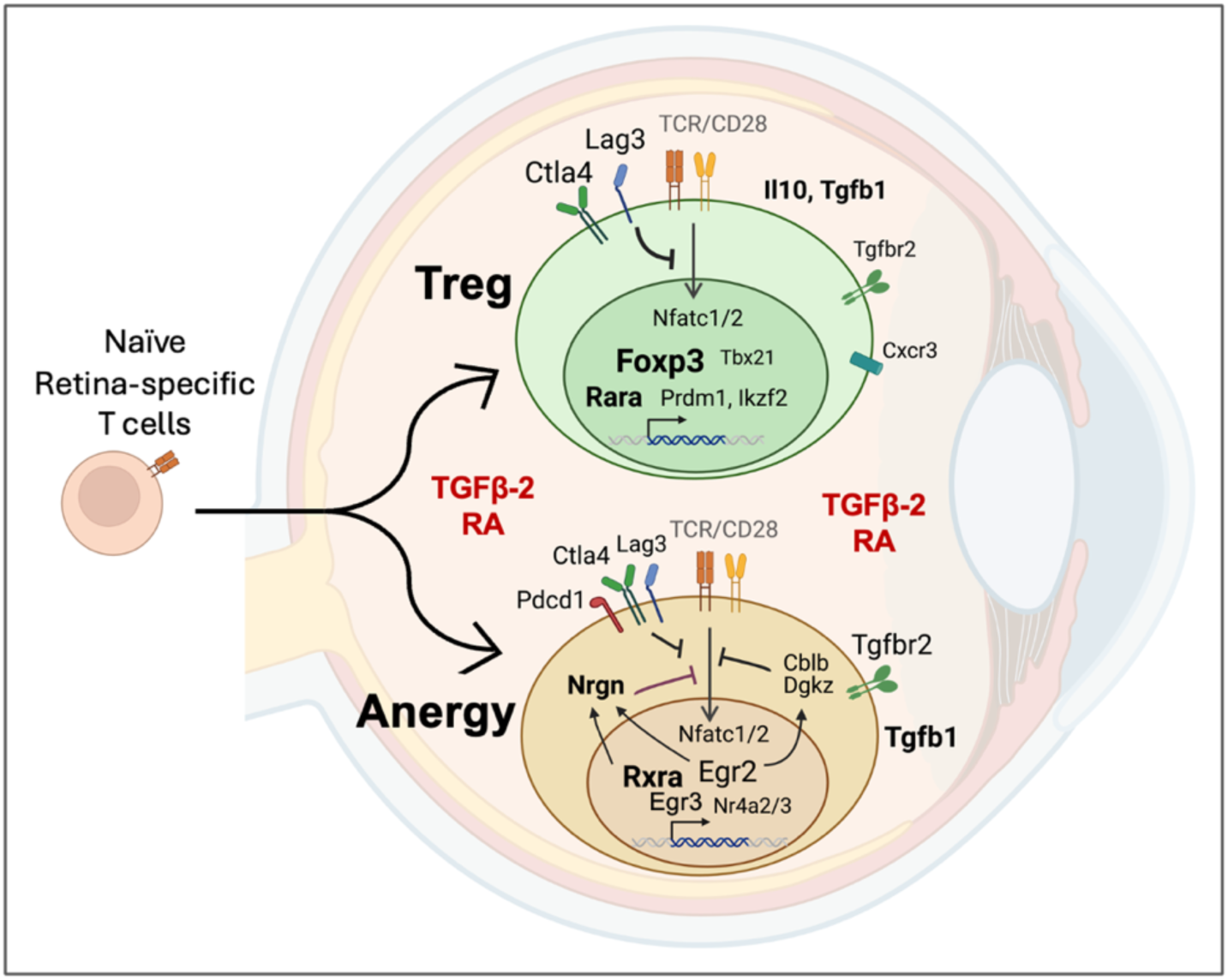

**Highlights:**

- Eye-primed retina-specific T cells develop distinct tolerance-associated phenotypes.
- Eye-induced anergic T cells remain hyporesponsive to antigen re-stimulation.
- Regulatory and anergic T cells differentiate in parallel from a common precursor.
- Induction of T cell anergy is a novel feature of ocular immune privilege.

## INTRODUCTION

The eye has developed evolutionary adaptations that limit local inflammation in order to protect vision, that collectively form the complex phenomenon known as ocular immune privilege ^1 2^. In addition to the physical blood-tissue barriers that separate the eye from the immune system, multiple studies described that the intraocular environment, composed of ocular fluids and ocular resident cells, is immunosuppressive and can inhibit the activity of immunocompetent cells ^3 4 5^. Aqueous humor (AH) has been shown to reduce proinflammatory cytokine production by T cells in culture and to promote induction of regulatory T cells (Tregs) ^6 7^. Soluble factors involved in these processes include transforming growth factor-beta (TGF-β), α-melanocyte-stimulating hormone (α-MSH), vasoactive intestinal peptide (VIP), retinoic acid (RA), and others ^5 2^. Retinal glial Müller cells were the first ocular resident cells shown to inhibit T cells in co-culture ^8^. Since then, many reports described induction of Tregs by pigmented epithelia in the front and back of the eye ^3 9^. Immunomodulatory molecules expressed by these cells include CDhi ^10^ and PD-LT ^11^ that engage the inhibitory receptors CTLA-E and PD-T on T cells, respectively, as well as membrane-bound or soluble TGF-β and CTLA-Yα ^12^. Retinal microglia and dendritic-like cells also have been reported to inhibit antigen-specific T cell responses and to induce Tregs, possibly through aberrant antigen presentation ^13 14 15^. However, these studies were conducted largely *in vitro*, and could not represent the complexity of the living eye. Further, many predated the discovery of Forkhead box PO (FoxpO) as a marker for Tregs, making it difficult to distinguish *de novo* induction of Tregs, from expansion of a preexisting Treg population.

Immune sequestration of unique retinal antigens (Ag), which are absent in the periphery, behind a blood-retinal barrier impedes development of peripheral tolerance in autoreactive T cells that escaped thymic negative selection ^2 16^. Such cells can be easily triggered to become pathogenic effectors, but nevertheless, autoimmune uveitis remains a relatively rare disease ^17^. To address the question how the eye maintains immune homeostasis, we established a mouse model in which retina-specific T cells, capable of inducing autoimmune uveitis, are injected into the eye ^18^. This model exposes the eye to naïve but non-tolerant T cells, as would occur in case of intraocular bleeding as a result of trauma or vascular abnormalities (e.g., macular degeneration, diabetic retinopathy, or neovascular glaucoma) ^19 20^. Interestingly, the T cells acquired an antigen-experienced (primed) phenotype within the eye but failed to induce uveitis. While ∼Ob% converted to FoxpO^+^ Tregs, the majority did not, and produced detectable levels of IFN-γ and IL-TUA ^18^. However, the fate and function of these eye-primed cells could not be determined, due to lack of appropriate technology.

In the current study, we utilized single-cell RNA sequencing (scRNA-seq) to comprehensively define the transcriptome of retina-specific T cells responding to their cognate antigen in the privileged intraocular environment. We present evidence that the non-FoxpO converted population is not effectors being kept in check by the Tregs, but rather represents a novel anergic phenotype unique to the eye that differentiates in parallel with FoxpO^+^ Tregs from naive retina-specific T cells. Our findings are the first to dissect the phenomenon of ocular immune privilege at the molecular level.

## RESULTS

### Naive retina-specific T cells differentiate into several distinct subtypes within the ocular environment

To gain better insight into the transcriptomic landscape that naive retina-specific T cells acquire within the eye, we performed scRNA-seq using the *in vivo* ocular immune privilege model. Briefly, naïve T cells were fluorescence-activated cell sorting (FACS)-sorted from *Tcra^−/−^* RTiTH FoxpO^GFP^ CDlb.Y mice and were intravitreally injected into the eyes of WT CDlb.T congenic recipients (Fig. 1A). The gating strategy for obtaining naive retina-specific non-Treg T cells from donors is depicted in Fig. 1B. One week after the injection, CDE^+^ CDlb.Y^+^ donor-derived T cells were retrieved from the eyes of CDlb.T congenic recipients. As we reported previously, about a third (OT.h%) of the cells converted to FoxpO^+^ phenotype (Fig. 1C), and represent functionally competent Tregs ^18^. The naive retina-specific T cells before intravitreal injection and the injected cells retrieved from the recipient’s eyes were labeled by hashtag oligos before pooling and subjected to scRNA-seq (Fig. 1A and Table S1).

**Fig. 1:**
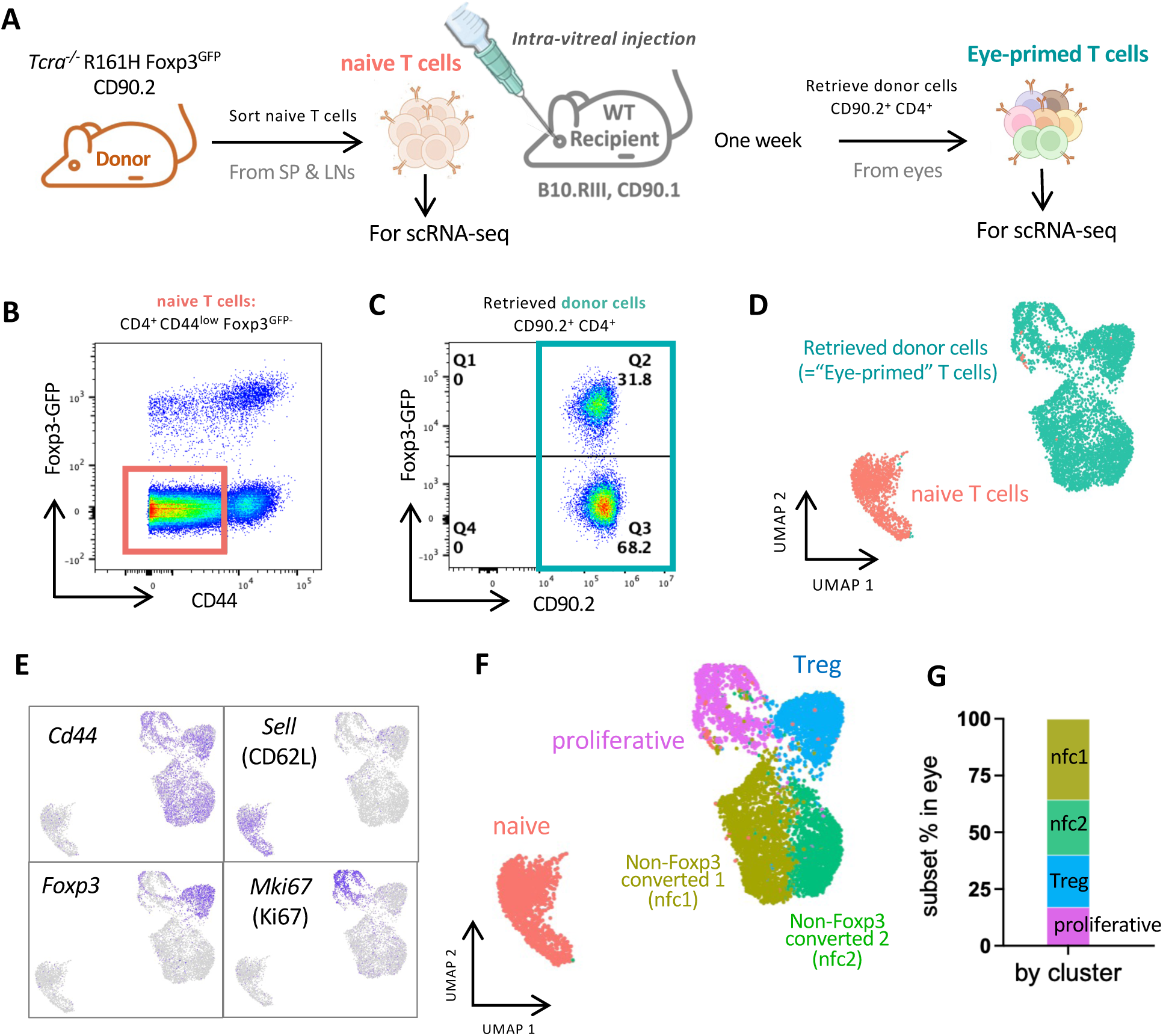
Naive retina-specific T cells differentiate into several distinct subtypes within the ocular environment. (A) Retina-specific naive T cells were sorted from peripheral lymphoid tissues (spleens and lymph nodes) of Tcra-/-R161H Foxp3GFP reporter donor mice and injected into the eyes of CD90.1 congenic WT recipients. One week later, the CD90.2 donor T cells (eye-primed T cells) were retrieved from both eyes of 4 recipients. Naive and eye-primed T cells were subjected to scRNA-seq. (B) Gating strategy for the naive donor cell sorting (CD4+CD44lowFoxp3GFP-). (C) CD90.2+ CD4+ T cells retrieved from eyes of recipients were analyzed for the Foxp3 expression by flow cytometry. (D) UMAP showing naive T cells and eye-primed T cells. (E) Feature plots showing expression of antigen-primed T cell phenotype (CD44+ CD62L-), Treg maker (Foxp3) and proliferation marker (Mki67) in eye-primed T cells indicated by purple color. (F) Cluster identification and (G) Ratios of T cell subtypes of eye-primed T cells.

Using standard scRNA-seq analysis pipelines ^21^, cells that passed quality control were used for downstream analyses. Cells before- and after-injection into the eyes showed a clear division in the UMAP (Fig. 1D). A distinct pattern of high *Cd44* and almost no CDiYL (*Sell*) expression was confirmed in all cells recovered from the eyes, indicating that they had been primed (Fig. 1E). The great majority of *Foxp3*^+^ Tregs formed one separate cluster, while a few *Foxp3^+^* Tregs were detected in the neighboring cluster with high levels of the proliferation marker *Mki67* (Fig. 1E). This was in line with our previous finding that acquisition of FoxpO expression in the ocular environment was accompanied by proliferation ^18^. Unsupervised clustering further uncovered five major populations (Fig. 1F) based on their differentially expressed genes (Figure S1A). Among them, naive (*Foxp1, Lef1, Satb1*), Treg (*Foxp3* and *Il10*), and proliferative (*Mki67, Stmn1*, *Top2a*) clusters were straightforward to annotate (Fig. 1E and Figure S1A). However, we did not find a clear pattern of gene expression to help classify the other two *Foxp3*-negative clusters as known lineages (Figure S1A). Therefore, for lack of a better definition, they are designated as “non-FoxpO converted clusters” (nfcT and nfcY) in the interim. The proportions of the T cell subpopulations recovered from recipients’ eyes are shown in Fig. 1G). Taken together, the scRNA-seq data revealed that the naive T cells exposed to their cognate antigen within the ocular environment differentiated from a homogeneous population into diverse subtypes, with Treg cells constituting one of several discrete populations.

### Non-FoxpK-converted (nfc) clusters do not conform to canonical gene patterns of known effector Th-lineages

To assess the phenotype of the non-FoxpO converted (nfc) cells, we screened defining gene sets, including master transcription factors (TFs), signature cytokines, chemokines and surface molecules characteristic of the known major T-effector lineages.

Levels of ThT and ThTU lineage-defining transcription factors, *Tbx21* for ThT and *Rorc* for ThTU, respectively, were relatively low across all four eye-primed clusters, and not restricted to any particular cluster (Fig. 2A-B). ThT-associated surface markers (*Ccr5* and *Cxcr3)* ^22^, were confined to the Treg cluster, but absent in the nfc clusters (Fig. 2B). Additionally, *Ifng, and Csf2*, key pro-inflammatory ThT cytokines, were undetectable in both nfc clusters (Fig. 2A), suggesting there is no ThT induction. Although a moderate *Il17a* was present in nfcY cluster (Fig. 2A), other ThTU signature genes (*Csf2, Il17f, Il22*, and *Il23r)* ^23,24^ were not detected (Fig. 2A and Figure S1B). The ThTU surface marker *Ccr6* was also not prominently expressed in nfcY (Fig. 2B), making it inconsistent with a canonical ThTU profile. The nfcT cluster exhibited modest expression of TF *Gata3* for ThY and *Bcl6* for T follicular helper (Tfh) cells (Fig. 2A), yet lacked other ThY-associated (*Ccr3, CcrL, Il4, Il5, IlM*) or Tfh-associated (*Cxcr5, Cxcl13, Il21*) genes (Figure S1B), suggesting that nfcT does not align with either ThY or Tfh lineage.

**Fig. 2:**
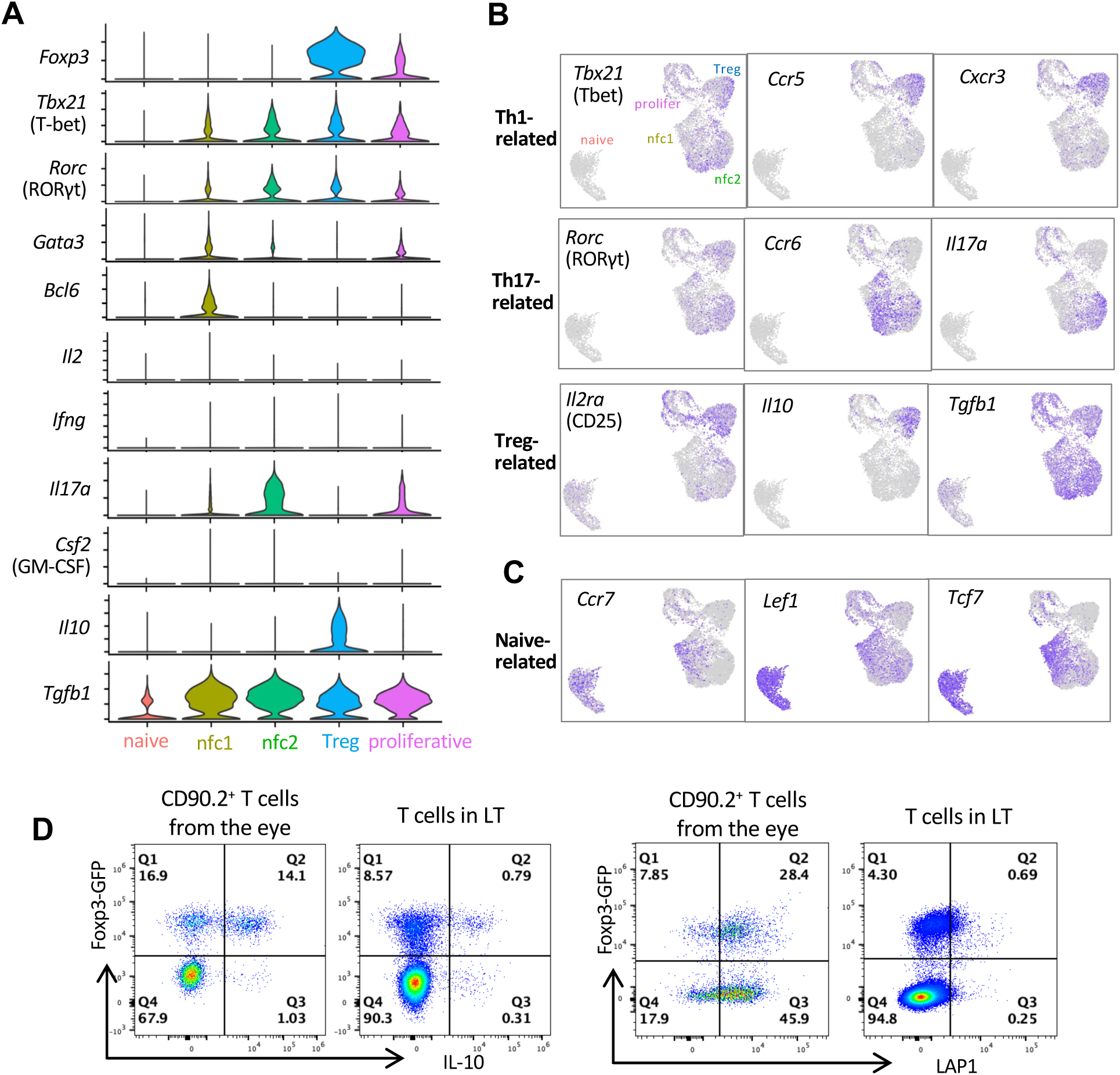
Non-Foxp3-converted (nfc) clusters do not conform to canonical gene patterns of effector Th-lineages. (A) Violin plots showing the frequency of cells expressing Th lineage-specific transcription factors (TFs) and cytokines in each cluster. (B) Feature plots showing the distribution of selected Th lineage-specific markers including TFs, cytokines and chemokine receptors. (C) Feature plots showing distribution of markers associated with a naïve-like phenotype (Ccr7, Lef1, Tcf7). (D) FACS plots showing LAP-1 (encoded by the Tgfb1 gene) and IL-10 expression in Foxp3-converted vs. non-Foxp3-converted recovered from eyes of recipient mice after 1 week. LT=lymphoid tissue T cells from naïve R161H donor mice.

Thl-related (*Spi1*, *Il4ra*, *IlM*) and ThYY-related (*Ahr*, *Ccr4*, *Ccl7, Il13*, *Il22*) genes were not observed either (Figure S1B), so these two lineages have no similarity with the nfc clusters. Interestingly, the nfcT and a part of the nfcY populations shared several T cell markers with naïve or resting cells (*Ccr7*, *Lef1*, and *Tcf7*) ^25^ (Fig. 2C).

We next considered FoxpO-negative regulatory phenotypes. Type T T-regulatory (TrT) cells are featured as IL-Tb producers independent of FoxpO^26^, but the nfc cells showed little to no *Il10* expression nor IL-Tb production (Fig. 2A-B and Fig. 2D), and this would argue against them being TrT cells. Instead, both nfc clusters showed high expression of *Tgfb1* (Fig. 2A-B), consistent with a ThO phenotype involved in mucosal immune regulation and oral tolerance ^27^. While they expressed LAP-T (latency-associated peptide, a product of *Tgfb1*), unlike some gut Tregs, they did not express IL-Tb ^28^ (Fig. 2D). Of note, *Tgfb2, Tgfb3* and the immunosuppressive cytokine IL-OV (*Il12a* and *Ebi3*) were undetectable in all eye-primed clusters (Figure S1C).

These data support the conclusion that the nfc clusters are Th-lineage-negative, and as such, are unlikely to represent known pathogenic effector cells or canonical Tregs.

### The nfc clusters exhibit a combination of regulatory/anergic gene signature

To better characterize the nfc clusters, we then investigated their global transcription profile. Gene signatures of each subset in the eye were defined by comparing their transcriptomes to that of naive T cells as baseline (Fig. 3A and Table S2). Both nfc clusters expressed high levels of anergy-associated genes as *Nrgn* ^29 30^, *Cblb* ^31^, *Dgkz* ^32^, *Nr4a1-3* ^33,34^, *Nrp1*, *Tox*, and *Tox2* ^16,35,36^. They also expressed canonical Treg-related genes, including inhibitory checkpoint molecules (*Nt5e*, *Maf*, *Itgav*, *Il2rb*, *Tnfrsf4*, *Tgfb1, Ctla4*, *Lag3*) ^37–41^ (Fig. 3A), and these were shared with the Treg cluster. Of note, *Ccl5*, *S100a4*, *S100a6*, *Tbx21*, and *Gzmb,* which characterize activated, highly suppressive Tregs ^38–40^ and were present in the Treg cluster, were not shared with the nfc populations (Fig. 3A).

**Fig. 3:**
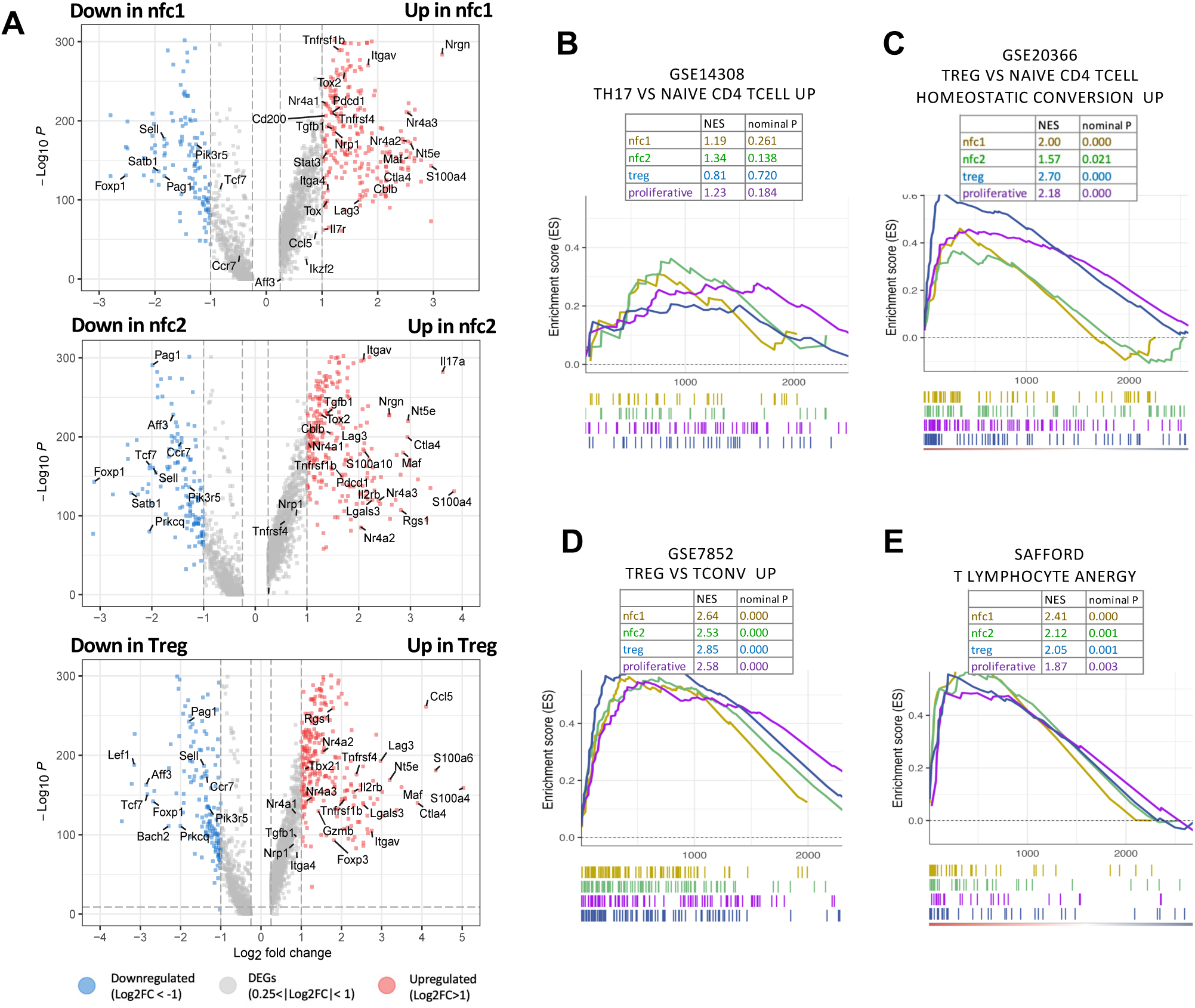
Intraocular environment induces regulatory and anergy signatures. (A) Volcano plots showing the differential gene expression of each cluster (baseline: naive cluster). Differentially expressed genes with more than 2-fold changes are highlighted in red (upregulated) or blue (downregulated). Representative signature genes are shown. (B-E) Representative GSEA enrichment results mapping the signatures of each cluster against the Molecular Signature Database (MSigDB). NES, normalized enrichment score, indicating the similarity of the current gene signatures with predefined gene sets. Nominal P value lower than 0.05 denotes significant similarity to the corresponding to the predefined gene set in B, C, D or E. (B) Signature of Th17 cells. (C) Signature of de novo converted Treg cells in vivo (also known as peripherally induced Tregs). (D) Signature of in vivo Treg cells in lymphoid tissues (spleen, thymus, and lymph nodes). (E) Curated anergy signature from canonical in vitro anergy-inducing conditions

We then performed Gene Set Enrichment Analysis (GSEA) to align the signatures of eye-primed T cell clusters with predefined gene sets in the Molecular Signature Database (MSigDB) ^42^ (Fig. 3B-E). In spite of upregulated *Il17a* (Fig. 3A, middle), the nfcY cluster did not have a characteristic ThTU signature ^43^ (Fig. 3B). Rather, the two nfc clusters appeared to share characteristics with Treg cells that had been *de novo* differentiated in non-ocular tissues *in vivo* ^37^ (Fig. 3C), Tregs isolated from lymphoid tissues of healthy mice ^44^ (Fig. 3D) and to *in vitro* induced anergic T cells ^45^ (Fig. 3E). The Treg cluster also shared genes with the anergic signature (Fig. 3E), supporting the notion that many phenotypic and mechanistic traits are shared between Treg and anergic T cells defined by other studies ^46–48^. Given that the nfc clusters lacked the defining Treg gene FoxpO, in the aggregate, they conformed best to the ‘T lymphocyte anergy’ gene set.

### The nfc clusters are hyporesponsive to antigenic stimulation but lack suppressive function

The results above suggested that the non-FoxpO-converted clusters were more reminiscent of Treg cells than of other Th cell lineages, and strongly resembled anergic T cells. Previous studies had demonstrated that a functional characteristic of anergy is hyporesponsiveness to antigenic stimulation, which can be rescued by IL-Y ^49^. To examine whether this was true of our nfc populations, we separated FoxpO+ and FoxpO– cells retrieved from the eye by flow sorting. Tregs and naïve T cells from peripheral lymphoid tissues (LT) of *Tcra^−/−^* RTiTH FoxpO^GFP^ transgenic mice were used for comparison. Proliferation was measured by [^3^H]-Thymidine incorporation in a co-culture system with antigen presenting cells (APC) and Interphotoreceptor Retinoid-Binding Protein (IRBP) peptide as cognate antigen (Ag) (Fig. 4A). Eye-induced non-FoxpO-converted cells exhibited minimal proliferation, which was considerably enhanced in the presence of IL-Y, although it remained markedly lower than that of the naïve cells (Fig. 4B). As expected ^50^, Treg cells also responded to IL-Y supplementation. Together with Fig 3, we interpret our data to mean that (a) the phenotype of the non-FoxpO converted T cells is consistent with anergy, and (b) these cells remain hyporesponsive to their cognate antigen independently of continued presence of FoxpO^+^ Tregs and of the ocular environment.

**Fig. 4:**
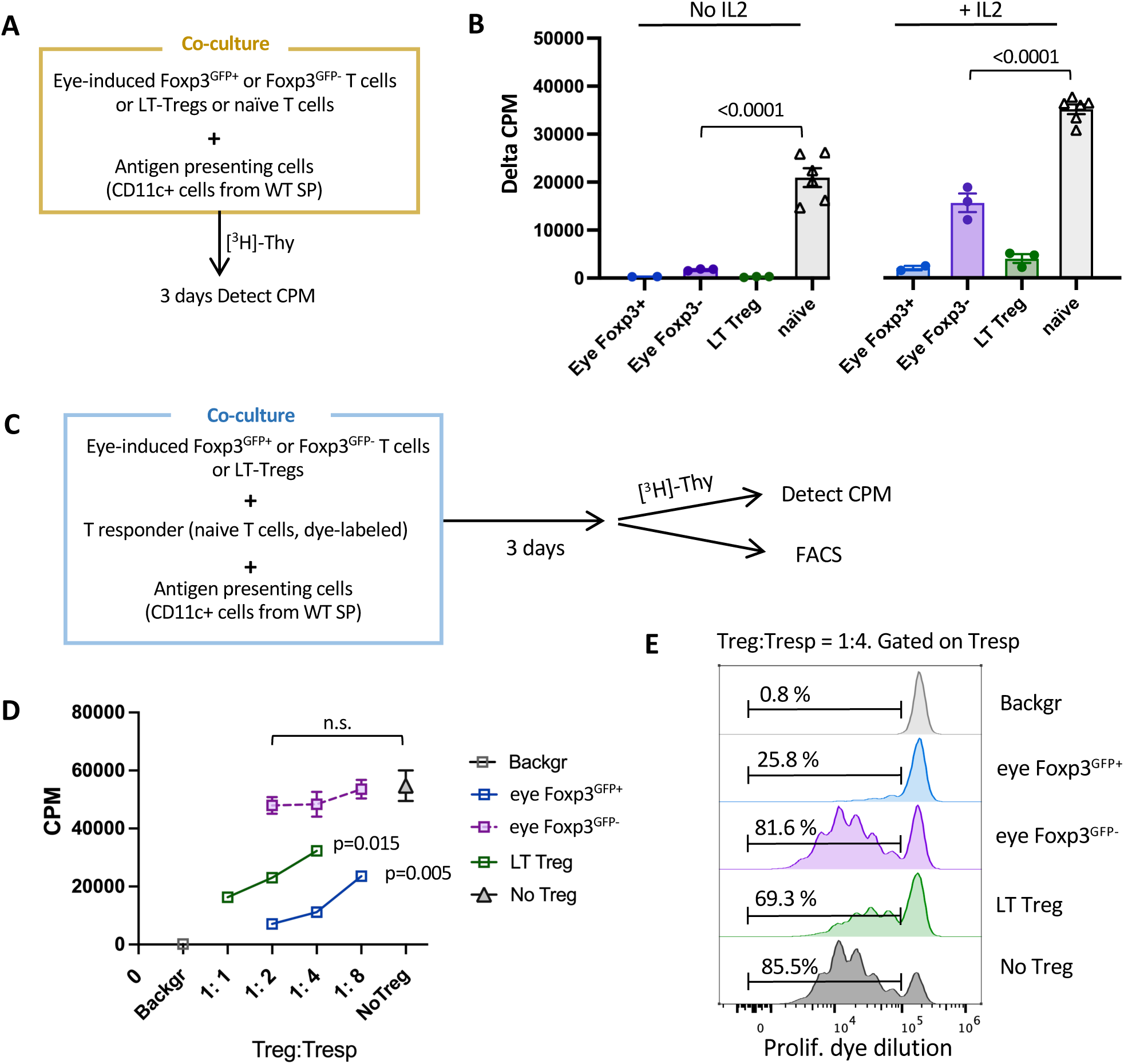
Non-Foxp3-converted (nfc) cells are functionally hyporesponsive to cognate antigen but not suppressive. (A) Diagram of co-culture system in B. Naïve T cells or Tregs from peripheral lymphoid tissues (LT) were sorted from Tcra-/-R161H Foxp3GFP mice. Donor-derived Tcra-/-R161H Foxp3GFP+ and Foxp3GFP– cells were sorted from recipients’ eyes. Antigen presenting CD11c+ cells were magnetically enriched from the spleen of WT B10.RIII mice. (B) Proliferation to the cognate antigen human IRBP161-180 in the presence or absence of IL-2. Cells without antigen served as background. Only one or two wells of eye-induced Treg cells could be set up per experiment. Shown is one representative experiment of 3. (C) Scheme for antigen-specific suppression assay by [3H]-Thymidine uptake and dye dilution methods for D and E. (D) Dose-dependent suppression of T responder cells by putative suppressors with antigen stimulation (triplicates). One representative experiment of two. (E) FACS plots of proliferation dye dilution. Tresp without antigen served as background.

Hyporesponsiveness to antigen that can be rescued by IL-Y is consistent with anergy, but is not necessarily a distinguishing attribute from Tregs. To resolve this, we compared the ability of eye-induced Treg and non-FoxpO-converted (FoxpO^GFP**–**^) T cells to suppress proliferation of naïve responders (Tresp) to their cognate Ag (IRBP peptide presented on splenic APC, Fig. 4C). We used two complementary methods: [^3^H]-Thymidine uptake and proliferation dye dilution, to exclude interference from possible proliferation of the Tregs themselves. Eye-induced Tregs and LT Tregs suppressed thymidine uptake by Tresp in a dose-dependent fashion. In contrast, the eye FoxpO^GFP**–**^ cells failed to significantly inhibit Tresp proliferation even at a T:Y Treg:Tresp ratio (Fig. 4D).

Dye dilution analysis confirmed that the proliferation rate of Tresp in the presence of anergic cells did not differ from control, indicating that FoxpO^GFP**–**^ cells lacked appreciable suppressor function *in vitro* (Fig. 4E).

Therefore, from here on, the non-FoxpO-converted nfcT and nfcY clusters will be referred to as ‘anergicT’ and ‘anergicY’, respectively.

### Distinct genes contribute to dampened TCR signaling in eye-induced anergic and regulatory cells

To address the question of what signaling molecules and pathways were involved in the induction the anergic and regulatory phenotypes, we performed an Ingenuity Pathway Analysis (IPA). Compared to the naïve cluster, pathways related to T cell suppression (T cell exhaustion, immunogenic cell death and TNFRY signals) and deficient T cell receptor (TCR) signaling (downregulated T cell receptor, CDYh, and ICOS signals) that restrain effector differentiation, were prominent in both Treg and anergic T cell clusters (Fig. 5A). Co-inhibitory genes that may feed into this pattern *Ctla4* and *Lag3* ^51,52^ were significantly upregulated in all eye-primed cell clusters, and their expression was highest in the Treg cluster, whereas *Pdcd1* (encoding PD-T), *Fasl* and *Cd200* were mainly increased in anergic clusters (Fig. 5B). Activation of TGF-β signaling in the anergic clusters (Fig. 5A) aligns with increased expression of TGF-β receptors (*Tgbr1–3*) (Fig. 5C), suggesting a connection between TGF-β signaling and the anergic state.

**Fig. 5:**
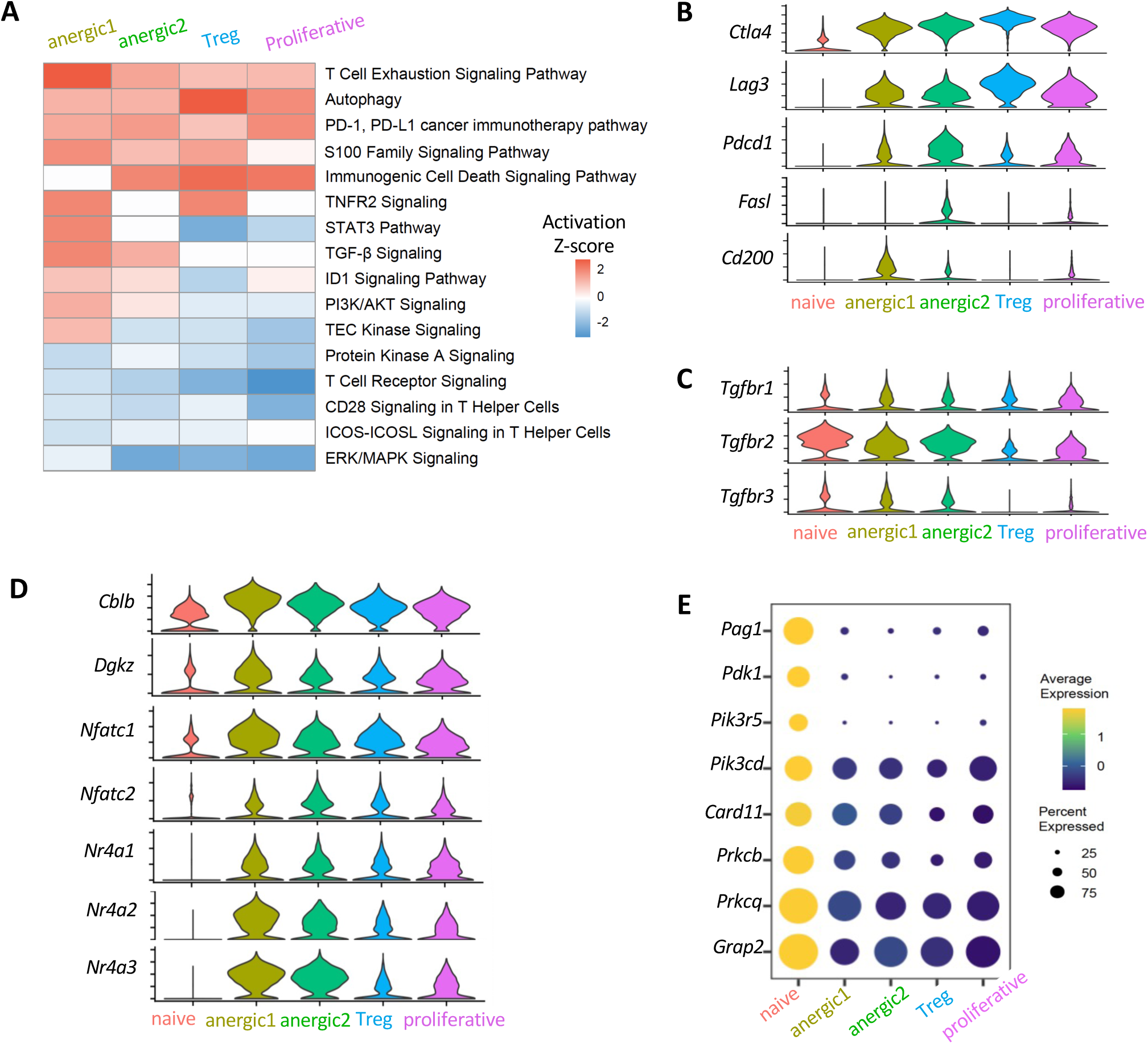
Suboptimal TCR signaling and inhibitory signals are involved in eye-induced tolerance. (A) Heatmap of differentially regulated immune-related signaling pathways in each T cell population compared to the naive baseline. Z score was calculated by Ingenuity Pathway Analysis (IPA). (B-C) Violin plots showing expression of canonical inhibitory markers (B), and TGF-β receptors (C). (D) Violin plots displaying gene expression of anergy-associated factors. (E) Bubble plot showing frequency of positive cells (size of bubble) and expression levels (color gradient) of genes involved in TCR/CD28 and MAPK signaling pathways..

Compared to Tregs, the anergic populations expressed higher levels of anergy-inducing factors such as NFAT (nuclear factor of activated T cells, encoded by *Nfatc1/2*) family members ^53 54^, NREA (nuclear receptor subfamily EA, encoded by *Nr4a1–3*) family members ^33,34^, CBL-B ^31^, and DGKζ ^32^, which are causally related to hyporesponsiveness, by affecting multiple components participating in TCR signal transduction ^16,55^ (Fig. 5D). This is consistent with the low expression of molecules downstream of TCR/CDYh pathways, such as pyruvate dehydrogenase kinase T (*Pdk1)*, PIOKs (*Pik3r5, Pik3cd*), and PKCs (*Prkcb, Prkcq*) (Fig. 5E), supporting dampened TCR/CDYh signaling.

### Distinct as well as shared tolerance-inducing regulons are enriched in anergic and Treg clusters

To identify potential key transcription factors (TF) regulating anergic and Treg subsets, we used the SCENIC (Single-Cell rEgulatory Network Inference and Clustering) pipeline, which infers TF activity from the co-expression of its direct target genes, collectively forming a regulon ^56^ (Fig. 6A). This provides a more precise readout of a cell’s state of differentiation than the TF mRNA expression alone.

**Fig. 6:**
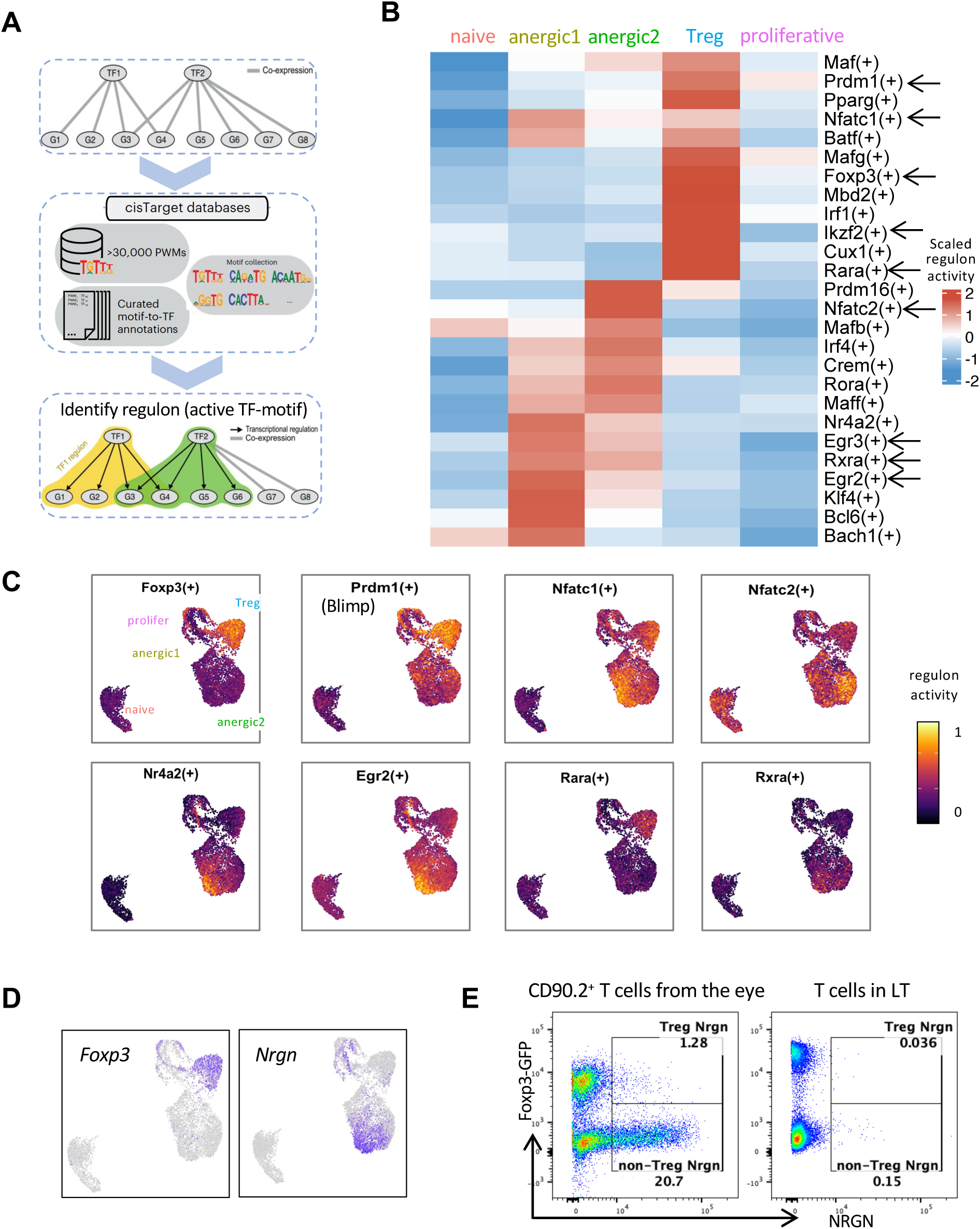
Distinct tolerance-inducing regulons and markers identify anergic and Treg cells. (A) Workflow of regulon identification by SCENIC (single-cell regulatory network inference and clustering) analysis. (B) Representative regulons enriched in eye-primed T cell subpopulations and naive T cells are marked with arrows in the heatmap. (C) Visualization of the regulon activity overlaid on the UMAP. (D and E) Reciprocal expression of Foxp3 and Nrgn in Treg and non-Treg (anergic) clusters by RNA (D) and protein (E).

Each of the anergic T clusters exhibited enrichment of distinct regulon activity. However, both clusters showed enrichment of regulons *Egr2*, *Egr3* ^45^, *Nr4a2,* and *Klf4* ^57^ (Fig. 6B). These TFs have been implicated in attenuating pathogenic T cell responses ^33^. Egr-Y in particular is considered a ‘master TF’ of anergy that directly upregulates CBL-B, DGKζ and other anergy-associated genes ^31,45,58^.

In the Treg cluster, FoxpO and its ‘accessory’ TFs *Prdm1* (encoding BlimpT), *Ikzf2* (Helios) and *Mbd2* (MbdY), that are known to promote and stabilize Treg function ^59–62^ exhibited increased regulon activities ^59^ (Fig. 6B). Enrichment of those FoxpO accessory regulons is compatible with the high suppressive function of the eye-induced Treg cells that we observed (Fig. 4D). A total of YVV regulons were identified as active within our dataset (Table S3). Of note, regulons corresponding to T-bet and RORγt were not enriched, confirming that eye-primed populations did not appear to be ThT or ThTU effectors.

Regulons active in both anergic and Treg clusters were *Nfatc1* and *Nfatc2* (Fig. 6C), which aligns with the known tolerogenic role of the NFAT family in both anergic cells and Tregs. In the former, they cooperate with NREA and EGRY/O, to repress effector cytokines (IFN-γ, GM-CSF) and to induce inhibitory regulators CTLA-E, PD-T, LAGO, CBL-B and DGKζ ^58,63^, and in the latter NFAT-FoxpO interaction upregulates Treg markers CTLA-E and CDYV, contributing to suppressor function ^54,64^. Moreover, we noticed the presence of regulons for the nuclear receptors for retinoic acid, *Rara and Rxra*. Of note, *Rara* regulon was highly activated in Tregs, whereas *Rxra* regulon was preferentially activated in anergic cells (Fig. 6B and 6C), suggesting that although RA regulates both Tregs and anergic cells, they rely on different receptors for RA mediated functions.

RARA is associated with FoxpO^+^ Treg development ^65^; however, since the anergic cluster lacks FoxpO, we looked for molecules downstream of the RXRA receptor. One gene known to be downstream of *Rxra*, *Nrgn* ^66^, was among the most highly expressed genes in the anergic clusters (Fig. 3A and Fig. 6D), and the top gene in the anergicT cluster. Notably, its expression closely matched the distribution of *Rxra* regulon activity (Fig. 6C-D) and was mutually exclusive with FoxpO expression (Fig. 6D). Control lymphoid tissue (LT) CDE^+^ cells, whether FoxpO^+^ or FoxpO^-^, lacked neurogranin (encoded by *Nrgn*) expression (Fig. 6E). *Nrgn* has normally been associated with neurons ^67^; here, we demonstrate high expression of *Nrgn* as well as its protein product, neurogranin, restricted to the eye-derived FoxpO-negative, i.e. anergic cells.

### Eye-induced Tregs and anergic cells seem to differentiate in parallel rather than sequentially

An important question in understanding the development of Treg and anergic cell fates is whether they differentiate as separate lineages or whether one derives from the other. To address this question, we performed trajectory analyses using RNA velocity ^68,69^ and Monocle pseudo-time trajectory analyses ^70^.

RNA velocity can infer the direction of cellular state changes and estimate the future state based on the relative abundance of spliced and un-spliced transcripts ^68^. After excluding the naïve population and reclustering the eye-primed cells, we projected the RNA velocity vectors onto a new UMAP (Fig. 7A). The trajectories originated from the proliferative cell cluster and diverged in distinct directions. The Treg and anergicT cluster appeared to be independent fates, with the arrows pointing in opposite directions, while a part of anergicY population appeared to transition toward the anergicT stage (Fig. 7A). Based on the estimated latent time by the scVelo algorithm ^69^, which reflects the internal clock of a cell (Fig. 7B), the anergicY cluster seems to represent a less differentiated state with lower latent time, whereas the Treg and anergicT populations may be more terminally differentiated.

**Fig. 7:**
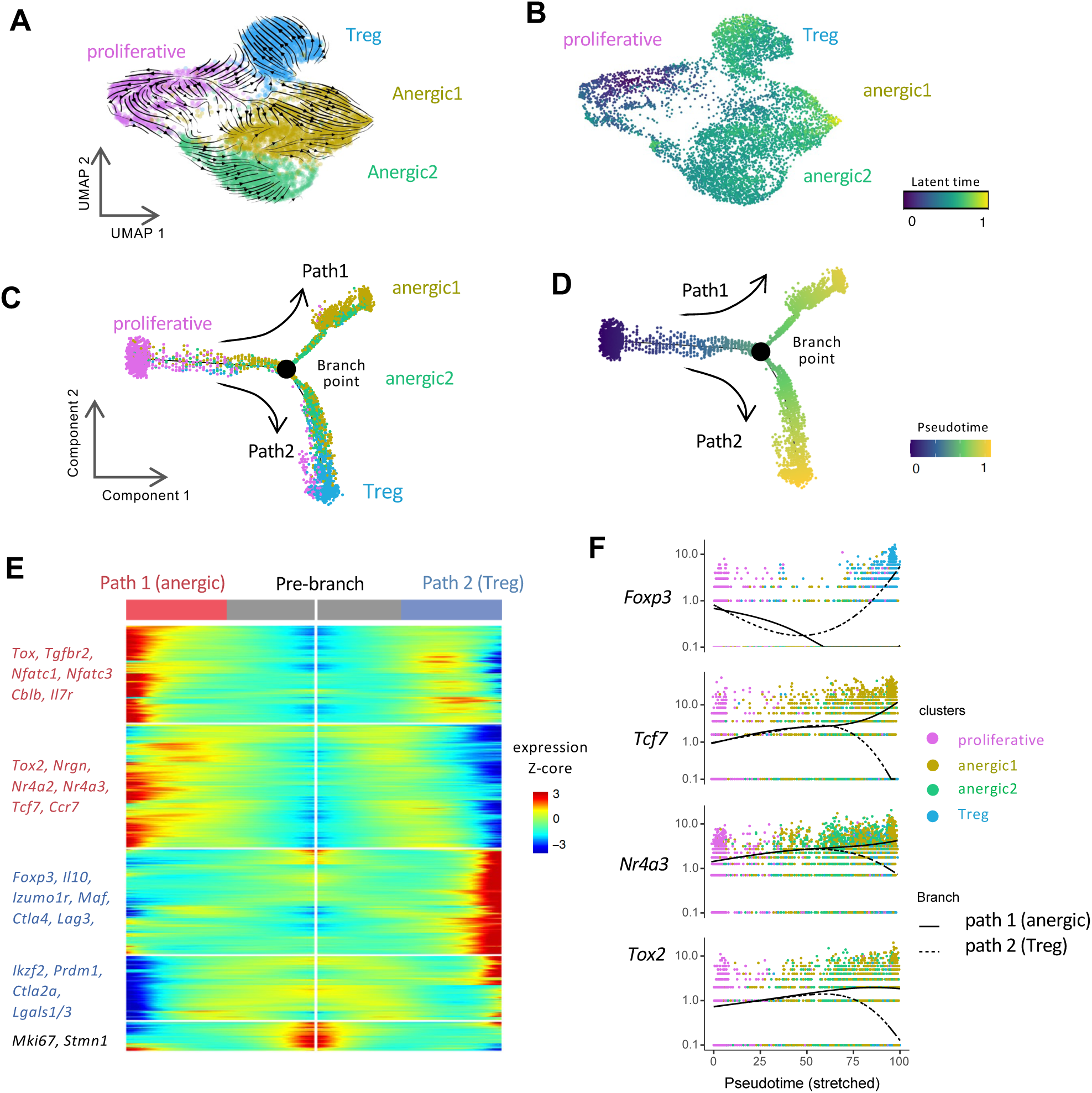
Trajectory analyses indicate parallel differentiation of Treg and anergic populations from an initial proliferative precursor. (A-B) RNA velocity analysis. Eye-primed cells were extracted and re-clustered for a new UMAP. (A) RNA velocity (arrowheads) projected onto this UMAP reflect the direction of cell state transitions. (B) The latent time is the calculated progression from the origin (proliferative cluster) to the end states (anergic1 and Treg), represented by color code. (C-F) Monocle pseudotime analysis. Monocle analysis ordered the cells along the pseudotime trajectories, displaying a branched pattern of two paths. (C) Color represents each cluster. (D) Color represents pseudotime, from the initial phase (0) to the late stage (1). (E) Heatmap showing the bifurcation of gene expression dynamics along pseudotime. (F) Kinetic patterns of specific canonical genes of the two paths. Cells were color-coded for each cluster.

As a complementary approach, we reconstructed the trajectories using the Monocle algorithm ^70^. The inferred state of cell transition revealed the emergence of two branches, arising from the common proliferative population and bifurcating at the branch point (Fig. 7C). Notably, one path was populated mainly by anergicT cells, and the other path was dominantly occupied by the Treg cells. The distribution of anergicY cells in both branches may point to the plasticity of this cluster (Fig. 7C). The branched pseudotime results supported that the anergicT and Treg clusters had reached their final stages (Fig. 7D).

Fig. 7E and Table S4 depict the genes undergoing the most pronounced dynamic changes when progressing along each of the two branched paths. Resting/quiescent state markers (*Ccr7, Il7r, Tcf7*) and anergy-associated genes (*Nfatc1/3, Nr4a2/3, Tox, Tox2, Cblb*) were progressively upregulated along the anergic path (Fig. 7E), suggesting that these cells gradually lost their effector potential. Conversely, higher levels of canonical Treg-associated genes (*Foxp3, Il10, Ikzf2, Prdm1*) along the Treg path, was consistent with progressive differentiation of Tregs (Fig. 7E). Furthermore, while representative anergy-associated genes, such as *Tcf7, Nr4a3,* and *Tox2*, progressively increased in the anergic path, they progressively decreased in the Treg path (Fig. 7F), recapitulating the dynamic acquisition of the respective phenotypes.

In summary, the trajectory analysis reveals a branched pattern in which naïve T cells primed within the eye differentiate largely in parallel, rather than in tandem, into Tregs and anergic T cells from a common proliferative precursor.

## DISCUSSION

We provide the first study that resolves at the single cell level how the living eye actively “disarms” the pathogenic potential of retina-specific T cells *in vivo*. Within the eye, incoming T cells encounter high levels of TGF-β (mainly the TGF-βY isoform) ^71^. Retinoic acid (RA) is also abundant in the eye, owing to its function in the visual cycle. This creates a unique environment that has a central role in ocular immune privilege ^18^. Outside the eye, RA is made by CDTbO^+^ DC in the gut, where it enhances Treg differentiation and may contribute to food tolerance ^72^. Within the eye, in addition to the FoxpO^+^ Treg fate adopted by a minority of the naïve T cells, we show that the remaining majority adopts a phenotype consistent with anergy. This finding fills a major gap in understanding that was left by our previous data ^18^, which found a dampened expression of effector cytokines and TFs at the population level, but could not distinguish effectors being kept in check by FoxpO^+^ Tregs, from an alternative cell fate(s), nor could it resolve possible subset(s). Our current data dissect this in detail at the molecular level, and resolve the cell fates and their differentiation trajectory.

### Anergy vs. Regulation: unique gene expression in ocular tolerance

The induction of anergic T cells in the living eye is a novel and little-explored aspect of ocular immune privilege. As mentioned in the Introduction, previous concepts of ocular immune privilege were based largely on *in vitro* studies with isolated cell populations or ocular fluids, and most of those studies dealt with induction of Tregs ^3 4 9^. Although one study suggested that interaction of T cells with RPE cells *in vitro* can result in anergy, for obvious reasons this does not reproduce the complexity of the actual intraocular environment ^9^. Moreover, the transcriptome of the affected T cells was not characterized. Our current study identifies many anergy-associated genes (*Ctla4, Lag3*, *Pdcd1, Cblb, Dgkz*), as well as activated anergy-promoting transcription factors (NFAT, EGRY/O, NREA, TOX families), that are shared with other models of T cell anergy ^16,55 73^. However, we also identify multiple genes whose expression pattern appears characteristic to eye-induced anergy.

Prominent examples are:

*(i) Nrgn* (neurogranin), which was in our hands restricted to the anergic T cell population, as was *Egr-2*, a known *Nrgn* inducer ^29^. Nrgn is constitutively expressed in neuronal cells, where it regulates synaptic plasticity ^67^. While a few studies reported *Nrgn* mRNA in lymphocytes ^22 30 29^, its functional contribution remains to be unraveled. The role of Nrgn is to modulate intracellular Ca++ levels through its interaction with Calmodulin ^74^. Specifically, Nrgn sequesters Calmodulin by physically binding to it, and makes it unavailable for binding with Ca++. RA promotes *Nrgn* gene expression by upregulating RA receptors, particularly RXR, which binds to the RA response elements (RARE) in the *Nrgn* promoter ^66,75^. Nrgn in turn binds to and sequesters Calmodulin, lowering available free calcium ^74^. We hypothesize that as this process occurs in the T cells that are in the process of differentiation in the eye, RA-driven RAR/RXR upregulation increases Nrgn, sequestering Calmodulin and reducing intracellular calcium and inhibiting Calcineurin and NFAT ^53 76^. Because Treg differentiation and functional activation requires high Ca++ levels ^53^, these conditions should skew the balance of Tregs and anergic T cells towards anergy. We propose that Nrgn, by regulating intracellular Ca^++^ availability, acts as a key checkpoint in the choice of anergic vs. regulatory cell fate by newly primed T cells differentiating from a common precursor in the TGF-β and RA-rich ocular environment. The validation of this central hypothesis in the regulation of ocular immune privilege and T cell anergy is the subject of a separate ongoing study.
*(ii)* Although anergic T cells are generally thought to lack cytokine expression, the ocular anergic cells expressed a high level of *Tgfb1. Tgfb1* was expressed also by ocular Tregs, and about Eb% of both populations expressed the TGF-β protein. To our knowledge, *Tgfb1* expression had not been previously reported in any model of anergic T cells, and may be a distinguishing feature of eye-induced anergy. Nevertheless, judging by the functional data, its expression did not confer regulatory function on the ocular anergic T cells.
*(iii)* An anergy-associated gene that was not significantly expressed in ocular anergic T cells is *Izumo1r* (encoding FRE), which, together with expression of *Nt5e* (encoding CDUO) and absence of FoxpO, is considered a defining phenotype of anergic T cells, but its function in anergy has not been elucidated. Our data suggest that it may not have a functional role in eye-induced anergy, or its role is redundant with that of a gene(s) differentially expressed in eye-derived vs. other anergic cells, such as *Dgkz, Cblb, Rgs1, Maf, Lgals*U, and *Furin* ^55^.

### Eye-induced anergic state differs from exhaustion

Although T cell anergy and exhaustion share many transcriptional features, the development of the eye-induced anergic T cells is inconsistent with exhaustion for several reasons. Exhaustion occurs in environments with strong antigenic stimulation and efficient antigen presentation ^77^. The healthy eye has few and quiescent professional antigen-presenting cells (APCs) ^78–80^. Inefficient antigen presentation is conducive to T cell anergy induction rather than exhaustion. As well, exhaustion typically requires chronic Ag stimulation and follows full activation for effector function, whereas the cells here were analyzed after only one week of Ag exposure, and the retina had minimal pathology ^18^. Finally, many exhaustion-associated genes, such as *Tigit, Havcr2, Shp1-2, Ptpn2, Blimp1* and *Irf4* ^77 35^ were undetectable or minimally expressed in eye-induced anergic cells.

### Effector Treg characteristics define the ocular Treg Population

The gene expression profile of eye-induced Tregs (*Il10*, *Tgfb1*, *Ctla4*, *Lag3*, and *Nt5e*) is consistent with a highly suppressive “effector Treg” phenotype ^38–41^. This was confirmed functionally by comparison with Tregs from spleen and lymph node tissues of the same animals (note that all T cells are IRBP-specific). Regulon analysis also uncovered that many “FoxpO accessory TFs”, such as BlimpT, Helios, and MbdY, are activated. These TFs help maintain Treg stability ^59–62^, suggesting that Tregs differentiated within the eye may have a stable phenotype. Of interest, eye-induced Tregs also displayed some ThT-like genes, as indicated by higher levels of *Tbx21*, *Cxcr3*, and *Ccr5* compared to non-FoxpO anergic cells, whereas the anergicY population shared the lineage-specific marker

*Il17a* with ThTU effector phenotype. Expression of lineage-specific genes shared with ThT and ThTU effector cells by Tregs is felt to facilitate interaction with the target effector population(s) ^81–83^. Uveitogenic effector T cells are a mixture of ThT and ThTU ^84^. It is therefore tempting to speculate that the ocular microenvironment diverts ‘would-be’ ThTU effectors to anergy, whereas ‘would-be’ ThT effectors are diverted to FoxpO^+^ Treg fate. Investigation of this hypothesis and of the unique eye-induced Treg phenotype is part of a separate ongoing study.

### Limitations of the study

While the *in vivo* model of immune privilege is a powerful tool to dissect eye-specific control of immune cell differentiation, the system also has limitations, both objective and subjective. The level of complexity of an *in vivo* system precludes analysis of the individual contributions of signals from each component that integrate to produce the final phenotypic and molecular events. In part, this could be addressed by including the various ocular resident cells in the analysis. By the same token, RNA-Seq performed at additional time points could provide further insights into the kinetics of the differentiation process that could have strengthened our conclusions from the trajectory analysis. However, technical and logistic difficulties inherent to this experimental model precluded addressing this in the current study.

### In conclusion

Our findings shed new light on the concepts of ocular immune privilege and the molecular mechanisms that actively maintain immunological homeostasis. The results lead to a model in which the ocular environment limits pathology by instructing the conversion of conventional T cells to Tregs or to an alternative fate of T cell anergy, rather than a scenario where a population of Tregs keeps a population of T effector cells in check. Identification of eye-induced regulatory and anergic signatures offers a valuable foundation for future research, and may inform therapeutic strategies for ocular inflammatory diseases. Furthermore, these unique signatures may inform strategies to reverse undesirable T cell unresponsiveness in contexts such as cancer, vaccination and chronic infection.

## MATERIALS AND METHODS

### Mice

Interphotoreceptor retinoid-binding protein (IRBP)-specific T cell receptor (TCR) transgenic (RTiTH), *Tcra* knockout mice (*Tcra^−/−^*) on the BTb.RIII background were generated as previously described ^85^ and were crossed to BTb.RIII FoxpO^GFP^ strain ^86^. *Tcra^−^*^/-^ RTiTH FoxpO^GFP^ reporter mice were used as donors for naive retina-specific T cells. CDlb.T congenic wildtype (WT) BTb.RIII mice were used as recipients. Both male and female mice i-Tb weeks old were used in this study. All animals were maintained under specific-pathogen-free conditions at NIH animal facility on standard chow and water ad libitum. Care and use of animals followed institutionally approved animal study protocols and Animal Research Advisory Committee (ARAC) guidelines.

### Ocular Immune Privilege Model

The ocular privilege model was established and described in our previous study ^18^. Briefly, retina-specific T cells, enriched from peripheral lymphoid tissues of *Tcra^−/−^* RTiTH FoxpO^GFP^ donor mice using CDO^+^ T cell enrichment columns (R&D Systems) or CDE^+^ T cell isolation kit (Miltenyi Biotec), were FACS sorted to obtain the naive population depleted of preexisting Tregs (CDEE^low^ CDYV^−^ FoxpO^GFP-^). WT CDlb.T-congenic recipient mice were injected intravitreally with Vbb,bbb of these naïve T cells in T.V microliters PBS into each eye, using a OOG needle and Hamilton syringe. The cells were retrieved from donor eyes U–h days later and prepared for analysis, as described ahead.

### Flow cytometry and cell sorting

Single-cell suspensions from spleens and lymph nodes (submandibular, axillary, inguinal, and mesenteric lymph nodes) collected from *Tcra^−/−^* RTiTH FoxpO^GFP^ CDlb.Y donor mice were used for isolation of retina-specific T cells. Cells were stained with surface antibodies and sorted for live CDE^+^ CDEE^low^ FoxpO^GFP-^ CDYV^−^ Dump^−^ cells to

∼ll% purity using FACSAria II and AriaIII/Fusion sorters (BD Biosciences). Non-CDE markers (CDh, NKT.T, BYYb, CDTTb, DXV, and GrT) were used for the dump channel. To retrieve retina-specific T cells from the eyes of CDlb.T congenic recipients, eyes were minced and treated with T mg/ml collagenase D for Ob min at OU°C. Donor-derived retina-specific T cells were sorted as live CDE^+^ CDlb.Y^+^ CDlb.T^−^ cells. Propidium Iodide or U-AAD (for sorting) and ViaKrome hbh (for flow analysis on CytoFlex LX, Beckman Coulter) were used to exclude dead cells. For intracellular staining of cytokine, cells were stimulated with PMA (Tb ng/ml) and ionomycin (Vbb ng/ml) in the presence of brefeldin A (GolgiPlug; BD) for E h, following staining for surface marker and live/dead cells. Cells were then fixed with E% paraformaldehyde for Ob mins and stained for intracellular proteins in Tx BD perm/wash buffer for T hour. For Nrgn staining, cells were stained with surface maker antibodies and then fixed with E% paraformaldehyde, followed by permeabilization and staining with Nrgn antibody at E°C for Ob mins, and AF-iEU conjugated anti-Rabbit secondary antibody for Yb mins. Antibodies used for cell sorting and flow cytometry analysis were from BD Biosciences, BioLegend, and eBioscience/ThermoFisher. Detailed antibody and clone information is listed in Table S1.

### Sample processing and scRNA-seq

Fresh naive donor T cells, or donor cells retrieved from recipient mouse eyes one week after intravitreal injection, were collected for scRNA-seq. Donor T cells in the eyes retrieved from each of the four recipient mice were individually labeled with anti-mouse TotalSeqB hashtags (BioLegend, Table S1). Individual samples were incubated with unique hashtags and sorting antibodies before FACS sorting, per manufacturer’s protocol. The viability of sorted cells was greater than lb%. Sorted single-cell suspensions were adjusted to Ubb–TYbb cells/μl before loading the TbX chromium chip. Samples were processed with the Chromium Next GEM Single Cell O’ reagent kit in the Chromium X platform following the standard protocol for O’ Gene Expression assay (TbX genomics). The gene expression and cell surface libraries were sequenced on NovaSeq ibbb platform (Illumina).

### Quality control and clustering of scRNA-seq data

Sequencing reads were demultiplexed and aligned using CellRanger (U.T.b) with the default parameters. The output matrix files were converted into a Seurat object for quality control and clustering. Standard scRNA-seq analysis (quality control, clustering, and marker gene detection) was performed using Seurat (vE.O.b) ^21^. Cells were excluded from analysis if they met any one of the following criteria: transcript counts less than Tbbb or more than Ebbbb, fewer than Ebb genes, more than h% mitochondrial fraction, ribosomal fraction less than Tb% or more than EV%. Highly variable features between individual cells were identified, and linear dimensional reduction was performed using principal component analysis (PCA). Unsupervised clusters were determined using the ‘FindNeighbors’ and ‘FindClusters’ functions based on the first Yb principal components (PCs). The clustering result was visualized using YD uniform manifold approximation and projection (UMAP). Clustering was done at b.Y resolution to keep the naive T cells as one “homogeneous” cluster. The ‘FindAllMarkers’ function was used to identify marker genes of each cluster within the data set.

### Signature identification and GSEA analysis

To characterize the phenotypes of donor T cells retrieved from recipient eyes, we defined their gene-expression signatures and performed Gene Set Enrichment Analysis (GSEA). The gene signatures were defined by comparing each cluster with the naive cluster using the ‘FindMarkers’ function. Genes were considered differentially expressed using the default Wilcoxon Rank Sum test and log fold-change (FC) threshold (Table S2). The full list of differentially expressed genes (DEGs) was ranked based on log FC and then mapped to the Molecular Signature Database (MSigDB) via GSEA software (vE.O.Y) ^42^.

### Antigen-specific proliferation assay

Ocular immune privilege model was conducted as described above, after one week, FoxpO^GFP+^ or FoxpO^GFP-^ CDE^+^ CDlb.Y^+^ CDlb.T^−^ cells were sorted out from the recipients’ eyes (Vb,bbb cells/ well) and co-cultured with human IRBP_161–180_ peptide (Vb ng/ml) and CDTTc^+^ dendritic cells (at a T:V ratio to T cells), with or without Tbb IU/ml recombinant human IL-Y. Dendritic cells were obtained by digesting spleens from WT CDlb.T mice in spleen dissociation medium (Stem Cell) for Ob minutes, followed by ammonium-chloride-potassium (ACK) lysis and CDTTc^+^ enrichment using Micro Beads (Miltenyi Biotec). Sorted naïve T cells (CDEE^low^ CDYV^−^ FoxpO^GFP-^ CDE^+^) and Treg cells (FoxpO^GFP+^ CDE^+^) from spleens and lymph nodes of *Tcra^−/−^* RTiTH FoxpO^GFP^ CDlb.Y mice were used as positive and negative controls, separately. Cell proliferation was determined using [^3^H]-Thymidine incorporation by adding TmCi/well after a Eh-hour culture and further incubated for Ti hours. Samples were harvested and counted using liquid scintillation (Perkin Elmer, MA). Unpaired Student *t*-tests were performed for statistics.

### Antigen-specific Treg suppression assays

Sorted FoxpO^GFP+^ or FoxpO^GFP-^ CDE^+^ CDlb.Y^+^ CDlb.T^−^ cells from recipients’ eyes were co-cultured with naïve retina-specific T cells (serve as T responder cells, Tresp; Vb,bbb cells/ well) and CDTTc^+^ dendritic cells (at a T:V ratio to Tresp). Treg cells (FoxpO^GFP+^ CDE^+^) from spleens and lymph nodes of *Tcra^−/−^* RTiTH FoxpO^GFP^ CDlb.Y mice were used as positive control of suppressor. Varying numbers of putative suppressor T cell populations were sorted and added to the cultures at the indicated Treg:Tresp ratios. Cell co-cultures were stimulated with Vbng/ml human IRBP_161-180_ without adding IL-Y. The inhibitory effect was assessed using either [^3^H]-Thymidine incorporation or proliferation dye dilution independently. For the dye dilution method, naïve T cells were labeled with proliferation dye - CellTracker DeepRed or CellTrace FarRed (Invitrogen/ThermoFisher Scientific) before setting up the culture. After O days, cells were stained for FACS analysis, and cell division of Tresp was quantified using FlowJo (Tb.h.b). Unpaired Student *t*-tests were performed for statistics.

### Ingenuity Pathway Analysis

Pathway analysis was performed using Ingenuity Pathway Analysis (IPA, www.qiagen.com/ingenuity). DEGs of each eye-primed cluster with corresponding log FC and adjusted P values were imported into IPA software for deciphering upregulated or downregulated functional pathways based on ingenuity knowledge base. After performing ‘core analysis’ of each T cell cluster independently, visualization across different clusters was achieved by ‘comparison analysis’ function. IPA’s z-score indicates a predicted activation or inhibition of a pathway, where a positive z value denotes an overall pathway’s activation and vice versa.

### The transcriptional factor activity (regulon) analysis

The python implementation of SCENIC (single-cell regulatory network inference and clustering, pySCENIC, vb.TY.T) ^56^ was used to predict the active transcriptional factor (TF). Starting from the normalized matrix data, the pySCENIC workflow consists of three stages. Initially, co-expression modules were inferred using a regression per-target approach. Then, the regulons (TF-target gene motifs) were refined from these modules based on cisTarget databases. Lastly, the ‘aucell’ algorithm was utilized to quantify the regulon activity score and find the significantly enriched regulon independently for each cell with default parameters. Information on software tools and cisTarget databases can be found in Table S3. The statistically significant regulons identified by SCENIC analysis were considered as active TFs, which reflected the upstream transcriptional drivers of the observed cellular identities ^56^. Regulon activity score was then scaled to plot heatmap or projected onto the UMAP.

### RNA velocity analysis

Velocyto ^68^ and scVelo ^69^ packages were used to perform RNA velocity analysis. First, RNA velocity (comprising spliced/un-spliced counts) for each cell was computed using the matrices generated by CellRanger and stored in the loom format. The velocity vectors were integrated into the Seurat object as a new data file. From the following file, we extracted the cells retrieved from recipient eyes and re-plotted the UMAP. The ‘latent time’ and ‘latent time facilitated RNA velocity’ were estimated using the likelihood-based dynamical model in scVelo. The velocity graph was visualized as streamlines overlaid by embedding in UMAP.

### Pseudotime analysis

Monocle (vY.Yi.b) ^70^ was applied to determine the potential lineage differentiation trajectory, keeping the default parameters. The matrix data of eye-primed clusters were imported as input for creating the Monocle ‘CellDataSet’. The ‘DDRTree’ method was utilized for dimensionality reduction and cell ordering along the pseudotime trajectory. To identify the genes that separate cells into branches, we performed the Branch Expression Analysis Modeling (BEAM) approach in Monocle Y. The dynamic expression of genes was visualized by the ‘plot_genes_branched_heatmap’ or ‘plot_genes_branched_pseudotime’ function.

## Abbreviations

Ag: Antigen
AH: Aqueous humor
APC: Antigen presenting cells
CBL-B: Casitas B-Lineage Lymphoma Proto-Oncogene B
CTLA-E: Cytotoxic T-Lymphocyte Associated Protein E
DEG: Differentially expressed genes
DGKζ: Diglyceride Kinase Zeta
EGR: Early Growth Response
FACS: Fluorescence-activated Cell Sorting
FC: Fold change
FoxpO: Forkhead box PO
GFP: Green fluorescent protein
GM-CSF: Granulocyte-Macrophage Colony-Stimulating Factor
GSEA: Gene Set Enrichment Analysis
ICOS: Inducible T-Cell Co-Stimulator
IFNγ: Interferon Gamma
IL-TUA: Interleukin TUA
IPA: Ingenuity Pathway Analysis
IRBP: Interphotoreceptor Retinoid-Binding Protein
LAP: Latency-Associated Peptide
LT: Lymphoid tissues
MAPK: Mitogen activated protein kinase
MsigDB: Molecular Signature Database
NFAT: Nuclear factor of activated T cells
Nrgn: Neurogranin
NREA: Nuclear receptor subfamily EA
PC: Principal component
PD-T: Programmed Cell Death T
PD-LT: Programmed Cell Death T Ligand T
PIOK: Phosphatidylinositol-E,V-Bisphosphate O-Kinase
PKC: Protein Kinase C
RA: Retinoic acid
RARA: Retinoic acid receptor alpha
RARE: Retinoic acid response elements
RXRA: Retinoid X receptor alpha
RPE: Retinal pigment epithelium
RORγt: Retinoid orphan receptor gamma t
SCENIC: Single-cell regulatory network inference and clustering
scRNA-seq: Single-cell RNA sequencing
TCR: T cell receptor
Th: T helper
TF: Transcription factor
Tfh: T follicular helper
TGF-β: Transforming growth factor-beta
TNFRY: Tumor Necrosis Factor Receptor Y
TOX: Thymocyte Selection-Associated High Mobility Group Box
TrT: TypeT regulatory cells
UMAP: Uniform manifold approximation and projection
VIP: Vasoactive intestinal peptide
WT: Wild type
α-MSH: α-melanocyte-stimulating hormon

## Data availability

The data reported in this paper are deposited in the Gene Expression Omnibus (GEO) database under accession no. GSEYhTETb. Code used for analysis can be found in GitHub https://github.com/NIH-NEI/Privilege_Treg_Anergy_scRNA.

## Funding

This study was supported by the National Eye Institute, National Institutes of Health (NIH) intramural funding (Project #EYbbbThE).

## Author contributions

Z.P.: Performed the experiments, analyzed the data, and wrote the manuscript draft. M.J.M and R.H: Planned the experiments, supervised the work, reviewed and edited the manuscript. V.N: Instructed and performed computational analyses, wrote software, and interpreted the results. Y.J: Developed methods and assisted in experiments. R.R.C: Conceptualized the study, acquired funding, reviewed, edited, and finalized the manuscript.

## Author disclosure statement

No competing financial interests exist.

## STAR METHODS

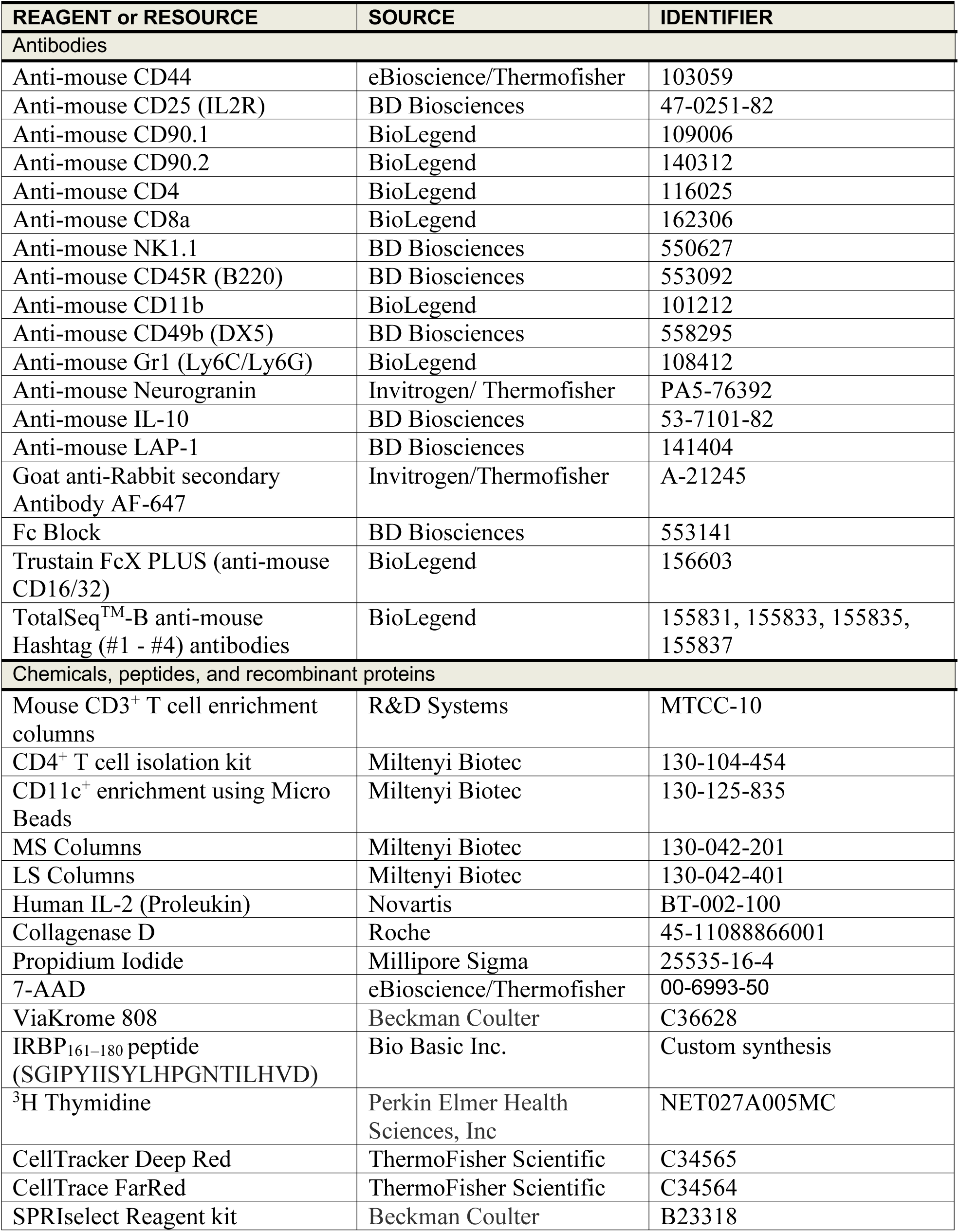
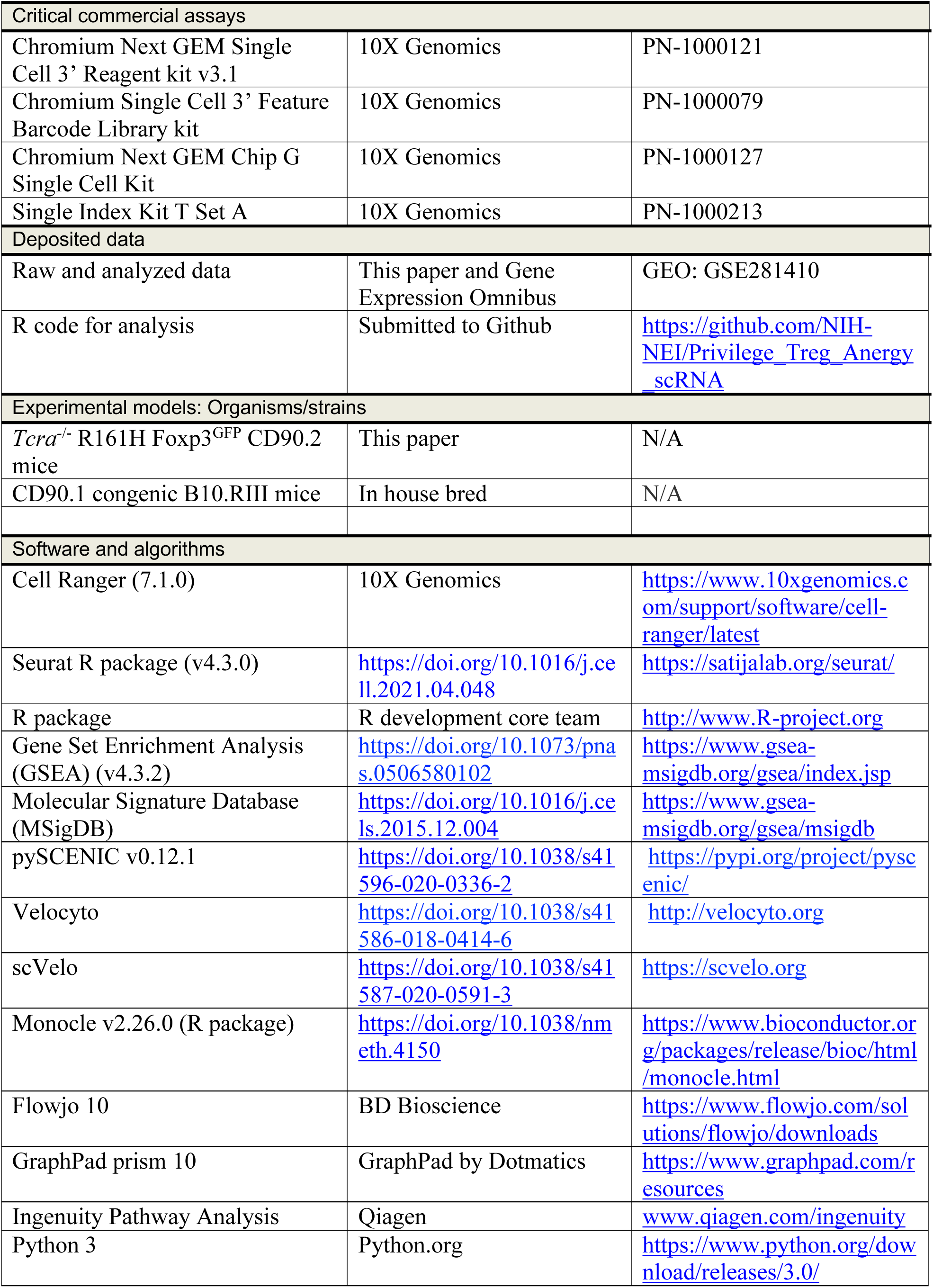
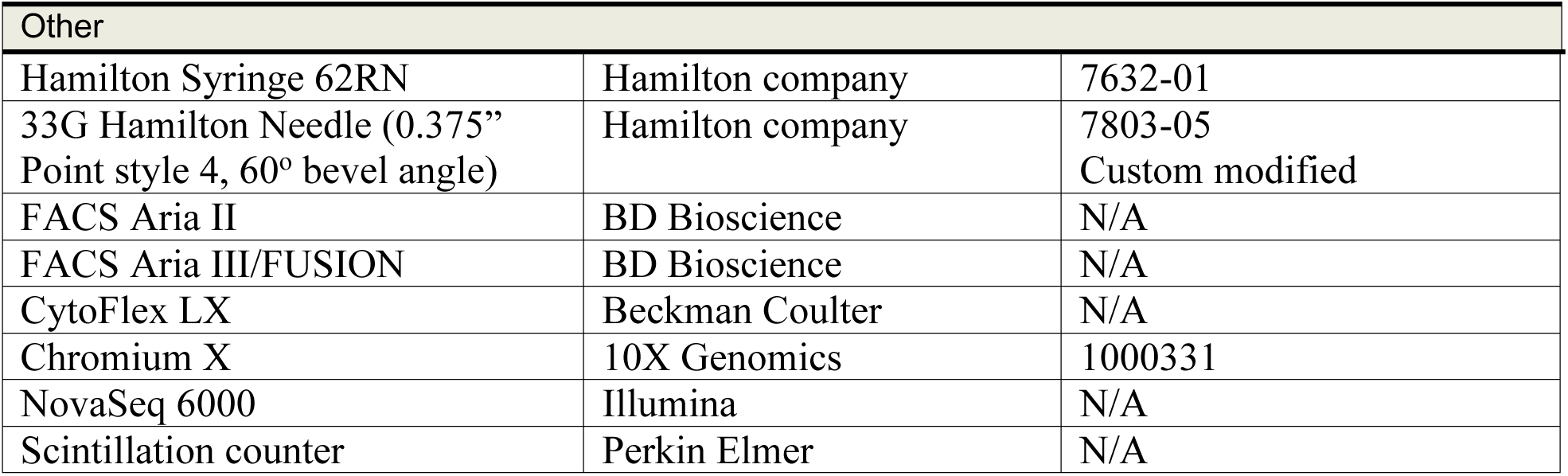

## Acknowledgements

The authors sincerely thank the National Eye Institute and National Heart Lung and Blood Institute Flow Cytometry Core facilities for assistance in conducting cell sorting, the Genetic Engineering Core Facility (NEI) for the generation of RTiTH TCR transgenic mice and genotyping service, and the National Cancer Institute CCR Genomics Core facility for sequencing. This study utilized the high-performance computational capabilities of the Biowulf Linux cluster at the NIH. We gratefully acknowledge the NIH Fellows Editorial Board for their valuable assistance in editing the manuscript for language and clarity. We thank Dr. Guangpu Shi (National Eye Institute, Laboratory of Immunology) for his comments and Dr. Han-Yu Shih for critically reviewing the manuscript. We are grateful to Drs. Nilisha Fernando and Jaanam Gopalakrishnan (National Eye Institute, Neuro-Immune Regulome Unit) for their assistance with scRNA-seq sample preparation. We would also like to express our gratitude to all the members of the Caspi Lab for their support and contributions.

## Appendix A. Supplementary data

Supplementary data to this article can be found online at https://doi.org/Tb.VYhT/zenodo.TEhiTEOb

---

**Figure. S1:**
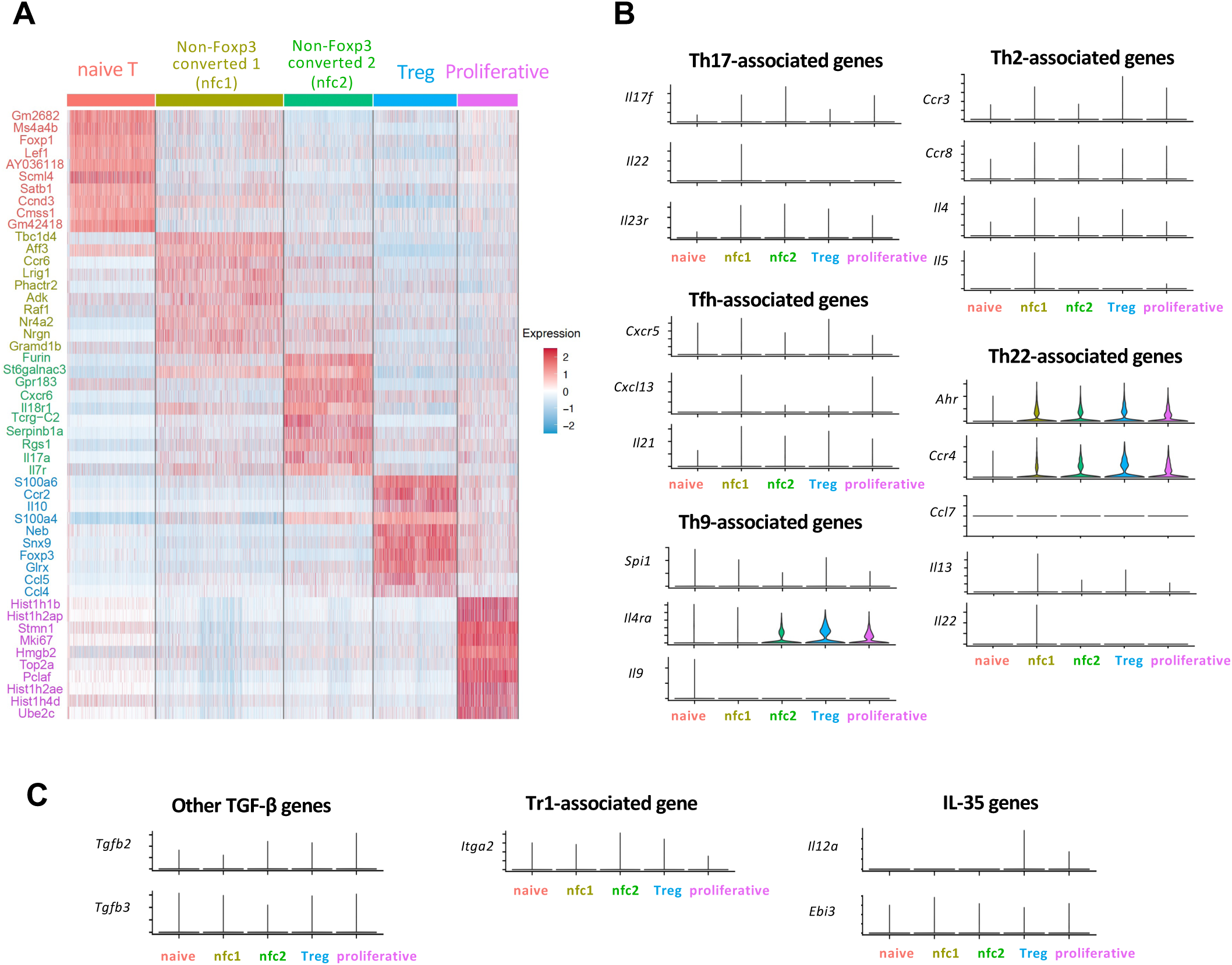
Transcriptional landscape suggests non-Foxp3-converted subsets are composed of Th-lineage-negative cells. (A) Heatmap of the top 10 differentially expressed genes in each cluster *vs.* all other clusters, showing scaled expression of the top 10 genes in each cluster. (B) Violin plots showing expression of Th effector lineage-related transcription factors (TFs), cytokines, chemokines/chemokine receptors: Th17-related (*Csf2, Il17f, Il22*, and *Il23r)*, Th2-associated (*Ccr3, Ccr8, Il4, Il5, Il9),* Tfh-associated (*Cxcr5, Cxcl13, Il21),* Th9-associated (*Spi1*, encoding PU.1, *Il4ra*, *Il9),* and Th22-associated (*Ahr*, *Ccr4*, *Ccl7*, *Il13*, *Il22*). (C) Violin plots showing the absence of Tr1-associated gene *Itga2* (encoding CD49b), and genes for TGF-β2, TGF-β3 and IL-35 (*Il12a, Ebi3*).

## Notes

https://doi.org/10.5281/zenodo.14861430

https://github.com/NIH-NEI/Privilege_Treg_Anergy_scRNA

### Competing Interest Statement

The authors have declared no competing interest.

## REFERENCES

1. Stein-Streilein, J., and Caspi, R.R. (2014). Immune privilege and the philosophy of immunology. Front Immunol 5,110. 10.3389/fimmu.2014.00110.

2. Taylor, A.W., and Ng, T.F. (2018). Negative regulators that mediate ocular immune privilege. J Leukocyte Biol 103, 1179–1187. 10.1002/Jlb.3mir0817-337r.

3. Mochizuki, M., Sugita, S., and Kamoi, K. (2013). Immunological homeostasis of the eye. Prog Retin Eye Res 33, 10–27. 10.1016/j.preteyeres.2012.10.002.

4. Gery, I., and Caspi, R.R. (2018). Tolerance Induction in Relation to the Eye. Front Immunol 9, 2304. 10.3389/fimmu.2018.02304.

5. Keino, H., Rorie, S., and Sugita, S. (2018). Immune Privilege and Eye-Derived T-Regulatory Cells. J Immunol Res 2018, 1679197. 10.1155/2018/1679197.

6. Taylor, A.W., Alard, P., Yee, D.G., and Streilein, J.W. (2007). Aqueous humor induces transforming growth factor-beta (TGF-beta)-producing regulatory T-cells. 1997. Ocul Immunol Inflamm 15, 215–224. 10.1080/09273940701382234.

7. Zhou, R., Horai, R., Mattapallil, M.J., and Caspi, R.R. (2011). A new look at immune privilege of the eye: dual role for the vision-related molecule retinoic acid. J Immunol 187, 4170–4177. 10.4049/jimmunol.1101634.

8. Caspi, R.R., Roberge, F.G., and Nussenblatt, R.B. (1987). Organ-resident, nonlymphoid cells suppress proliferation of autoimmune T-helper lymphocytes. Science 237, 1029–1032. 10.1126/science.2956685.

9. Gregerson, D.S., Reuss, N.D., Lew, K.L., McPherson, S.W., and Ferrington, D.A. (2007). Interaction of retinal pigmented epithelial cells and CD4 T cells leads to T-cell anergy. Invest Ophthalmol Vis Sci 48, 4654–4663. 10.1167/iovs.07-0286.

10. Usui, Y., Okunuki, Y., Hattori, T., Kezuka, T., Keino, H., Ebihara, N., Sugita, S., Usui, M., Goto, H., and Takeuchi, M. (2008). Functional expression of B7Hl on retinal pigment epithelial cells. Exp Eye Res 86, 52–59. 10.1016/j.exer.2007.09.007.

11. Sugita, S., Usui, Y., Rorie, S., Futagami, Y., Aburatani, H., Okazaki, T., Honjo, T., Takeuchi, M., and Mochizuki, M. (2009). T-cell suppression by programmed cell death l ligand 1 on retinal pigment epithelium during inflammatory conditions. Invest Ophthalmol Vis Sci 50, 2862–2870. 10.1167/iovs.08-2846.

12. Sugita, S., Rorie, S., Nakamura, O., Futagami, Y., Takase, H., Keino, H., Aburatani, H., Katunuma, N., Ishidoh, K., Yamamoto, Y., and Mochizuki, M. (2008). Retinal pigment epithelium-derived CTLA-2alpha induces TGFbeta-producing T regulatory cells. J Immunol 181, 7525–7536. 10.4049/jimmunol.181.11.7525.

13. Hsu, S.M., Mathew, R., Taylor, A.W., and Stein-Streilein, J. (2014). Ex-vivo tolerogenic F4/80(+) antigen-presenting cells (APC) induce efferent CD8(+) regulatory T cell-dependent suppression of experimental autoimmune uveitis. Clin Exp Immunol 176, 37–48. 10.1111/cei.12243.

14. Reuss, N.D., Lehmann, U., Norbury, C.C., McPherson, S.W., and Gregerson, D.S. (2012). Local activation of dendritic cells alters the pathogenesis of autoimmune disease in the retina. J Immunol 188, 1191–1200. 10.4049/jimmunol.1101621.

15. McPherson, S.W., Reuss, N.D., Pierson, M.J., and Gregerson, D.S. (2014). Retinal antigen-specific regulatory T cells protect against spontaneous and induced autoimmunity and require local dendritic cells. J Neuroinflammation 11, 205. 10.1186/sl2974-014-0205-4.

16. ElTanbouly, M.A., and Noelle, R.J. (2021). Rethinking peripheral T cell tolerance: checkpoints across a T cell’s journey. Nat Rev Immunol 21, 257–267. 10.1038/s41577-020-00454-2.

17. Garcia-Aparicio, A., Garcia de Yebenes, M.J., Oton, T., and Munoz-Fernandez, S. (2021). Prevalence and Incidence of Uveitis: A Systematic Review and Meta-analysis. Ophthalmic Epidemiol 28, 461–468. 10.1080/09286586.2021.1882506.

18. Zhou, R., Horai, R., Silver, P.B., Mattapallil, M.J., Zarate-Blades, C.R., Chong, W.P., Chen, J., Rigden, R.C., Villasmil, R., and Caspi, R.R. (2012). The living eye “disarms” uncommitted autoreactive T cells by converting them to Foxp3(+) regulatory cells following local antigen recognition. J Immunol 188, 1742–1750. 10.4049/jimmunol.1102415.

19. Rein, D.B., Wittenborn, J.S., Burke-Conte, Z., Gulia, R., Robalik, T., Ehrlich, J.R., Lundeen, E.A., and Flaxman, A.D. (2022). Prevalence of Age-Related Macular Degeneration in the US in 2019. JAMA Ophthalmol 140, 1202–1208. 10.1001/jamaophthalmol.2022.4401.

20. Choovuthayakorn, J., Worakriangkrai, V., Patikulsila, D., Watanachai, N., Kunavisarut, P., Chaikitmongkol, V., Luewattananont, D., and Tananuvat, N. (2020). Epidemiology of Eye Injuries Resulting in Hospitalization, a Referral Hospital-Based Study. Clin Ophthalmol 14, 1–6. 10.2147/OPTH.S234035.

21. Butler, A., Hoffman, P., Smibert, P., Papalexi, E., and Satija, R. (2018). Integrating single-cell transcriptomic data across different conditions, technologies, and species. Nat Biotechnol 36, 411-+. 10.1038/nbt.4096.

22. Kiner, E., Willie, E., Vijaykumar, B., Chowdhary, K., Schmutz, H., Chandler, J., Schnell, A., Thakore, P.I., LeGros, G., Mostafavi, S., et al. (2021). Gut CD4(+) T cell phenotypes are a continuum molded by microbes, not by T(H) archetypes. Nat Immunol 22, 216–228. 10.1038/s41590-020-00836-7.

23. Lee, Y., Awasthi, A., Yosef, N., Quintana, F.J., Xiao, S., Peters, A., Wu, C., Kleinewietfeld, M., Kunder, S., Hafler, D.A., et al. (2012). Induction and molecular signature of pathogenic TH17 cells. Nat Immunol 13, 991–999. 10.1038/ni.2416.

24. van der Veeken, J., Campbell, C., Pritykin, Y., Schizas, M., Verter, J., Hu, W., Wang, Z.M., Matheis, F., Mucida, D., Charbonnier, L.M., et al. (2022). Genetic tracing reveals transcription factor Foxp3-dependent and Foxp3-independent functionality of peripherally induced Treg cells. Immunity 55, 1173-+. 10.1016/j.immuni.2022.05.010.

25. Elyahu, Y., Hekselman, I., Eizenberg-Magar, I., Berner, O., Strominger, I., Schiller, M., Mittal, K., Nemirovsky, A., Eremenko, E., Vital, A., et al. (2019). Aging promotes reorganization of the CD4 T cell landscape toward extreme regulatory and effector phenotypes. Sci Adv 5, eaaw8330. 10.1126/sciadv.aaw8330.

26. Roncarolo, M.G., Gregori, S., Bacchetta, R., Battaglia, M., and Gagliani, N. (2018). The Biology of T Regulatory Type I Cells and Their Therapeutic Application in Immune-Mediated Diseases. Immunity 49, 1004–1019. 10.1016/j.immuni.2018.12.001.

27. Ochi, H., Abraham, M., Ishikawa, H., Frenkel, D., Yang, K., Basso, A.S., Wu, H., Chen, M.L., Gandhi, R., Miller, A., et al. (2006). Oral CD3-specific antibody suppresses autoimmune encephalomyelitis by inducing CD4+ CD25-LAP+ T cells. Nat Med 12, 627–635. 10.1038/nml408.

28. Branchett, W.J., Saraiva, M., and O’Garra, A. (2024). Regulation of inflammation by Interleukin-IO in the intestinal and respiratory mucosa. Curr Opin Immunol 91, 102495. 10.1016/j.coi.2024.102495.

29. Zheng, Y., Zha, Y.Y., Spaapen, R.M., Mathew, R., Barr, K., Bendelac, A., and Gajewski, T.F. (2013). Egr2-dependent gene expression profiling and ChIP-Seq reveal novel biologic targets in T cell anergy. Mol Immunol 55, 283–291. 10.1016/j.molimm.2013.03.006.

30. Merrell, K.T., Benschop, R.J., Gauld, S.B., Aviszus, K., Decote-Ricardo, D., Wysocki, L.J., and Cambier, J.C. (2006). Identification of anergic B cells within a wild-type repertoire. Immunity 25, 953–962. 10.1016/j.immuni.2006.10.017.

31. Nguyen, T.T.T., Wang, Z.E., Shen, L., Schroeder, A., Eckalbar, W., and Weiss, A. (2021). Cbl-b deficiency prevents functional but not phenotypic T cell anergy. J Exp Med 218. 10.1084/jem.20202477.

32. Olenchock, B.A., Guo, R., Carpenter, J.H., Jordan, M., Topham, M.K., Koretzky, G.A., and Zhong, X.P. (2006). Disruption of diacylglycerol metabolism impairs the induction of T cell anergy. Nature Immunology 7, 1174–1181. 10.1038/nil400.

33. Hiwa, R., Nielsen, H.V., Mueller, J.L., Mandla, R., and Zikherman, J. (2021). NR4A family members regulate T cell tolerance to preserve immune homeostasis and suppress autoimmunity. JCI Insight 6. 10.1172/jci.insight.151005.

34. Liu, X.D., Wang, Y., Lu, H.P., Li, J., Yan, X.W., Xiao, M.L., Hao, J., Alekseev, A., Khong, H., Chen, T.H., et al. (2019). Genome-wide analysis identifies NR4Al as a key mediator of T cell dysfunction. Nature 567, 525-+. 10.1038/s41586-019-0979-8.

35. Seo, H., Chen, J., Gonzalez-Avalos, E., Samaniego-Castruita, D., Das, A., Wang, Y.H., Lopez-Moyado, I.F., Georges, R.O., Zhang, W., Onodera, A., et al. (2019). TOX and TOX2 transcription factors cooperate with NR4A transcription factors to impose CD8(+) T cell exhaustion. Proc Natl Acad Sci US A 116, 12410–12415. 10.1073/pnas.1905675116.

36. Trefzer, A., Kadam, P., Wang, S.H., Pennavaria, S., Lober, B., Akcabozan, B., Kranich, J., Brocker, T., Nakano, N., Irmler, M., et al. (2021). Dynamic adoption of anergy by antigen-exhausted CD4(+) T cells. Cell Rep 34, 108748. 10.1016/j.celrep.2021.108748.

37. Feuerer, M., Hill, J.A., Kretschmer, K., von Boehmer, H., Mathis, D., and Benoist, C. (2010). Genomic definition of multiple ex vivo regulatory T cell subphenotypes. Proc Natl Acad Sci US A 107, 5919–5924. 10.1073/pnas.1002006107.

38. Dikiy, S., and Rudensky, A.Y. (2023). Principles of regulatory T cell function. Immunity 56, 240–255. 10.1016/j.immuni.2023.01.004.

39. Miragaia, R.J., Gomes, T., Chomka, A., Jardine, L., Riedel, A., Hegazy, A.N., Whibley, N., Tucci, A., Chen, X., Lindeman, I., et al. (2019). Single-Cell Transcriptomics of Regulatory T Cells Reveals Trajectories of Tissue Adaptation. Immunity 50, 493–504 e497. 10.1016/j.immuni.2019.01.001.

40. Scheinecker, C., Goschl, L., and Bonelli, M. (2020). Treg cells in health and autoimmune diseases: New insights from single cell analysis. J Autoimmun 110, 102376. 10.1016/j.jaut.2019.102376.

41. Zemmour, D., Zilionis, R., Kiner, E., Klein, A.M., Mathis, D., and Benoist, C. (2018). Single-cell gene expression reveals a landscape of regulatory T cell phenotypes shaped by the TCR. Nat Immunol 19, 291–301. 10.1038/s41590-018-0051-0.

42. Liberzon, A., Birger, C., Thorvaldsdottir, H., Ghandi, M., Mesirov, J.P., and Tamayo, P. (2015). The Molecular Signatures Database (MSigDB) hallmark gene set collection. Cell Syst 1, 417–425. 10.1016/j.cels.2015.12.004.

43. Wei, G., Wei, L., Zhu, J., Zang, C., Hu-Li, J., Yao, Z., Cui, K., Kanno, Y., Roh, T.Y., Watford, W.T., et al. (2009). Global mapping of H3K4me3 and H3K27me3 reveals specificity and plasticity in lineage fate determination of differentiating CD4+ T cells. Immunity 30, 155–167. 10.1016/j.immuni.2008.12.009.

44. Feuerer, M., Herrero, L., Cipolletta, D., Naaz, A., Wong, J., Nayer, A., Lee, J., Goldfine, A.B., Benoist, C., Shoelson, S., and Mathis, D. (2009). Lean, but not obese, fat is enriched for a unique population of regulatory T cells that affect metabolic parameters. Nat Med 15, 930–939. 10.1038/nm.2002.

45. Safford, M., Collins, S., Lutz, M.A., Allen, A., Huang, C.T., Kowalski, J., Blackford, A., Horton, M.R., Drake, C., Schwartz, R.H., and Powell, J.D. (2005). Egr-2 and Egr-3 are negative regulators of T cell activation (vol 6, pg 472, 2005). Nature Immunology 6, 737–737. 10.1038/ni0705-737.

46. Kalekar, L.A., and Mueller, D.L. (2017). Relationship between CD4 Regulatory T Cells and Anergy In Vivo. Journal of Immunology 198, 2527–2533. 10.4049/jimmunol.1602031.

47. Gavin, M.A., Clarke, S.R., Negrou, E., Gallegos, A., and Rudensky, A. (2002). Homeostasis and anergy of CD4(+)CD25(+) suppressor T cells in vivo. Nat Immunol 3, 33–41. 10.1038/ni743.

48. Maggi, J., Schafer, C., Ubilla-Olguín, G., Catalán, D., Schinnerling, K., and Aguillón, J.C. (2015). Therapeutic Potential of Hyporesponsive CD4(+) T Cells in Autoimmunity. Front Immunol 6,488. 10.3389/fimmu.2015.00488.

49. ElTanbouly, M.A., and Noelle, R.J. (2021). Rethinking peripheral T cell tolerance: checkpoints across a T cell’s journey. Nature Reviews Immunology 21, 257–267. 10.1038/s41577-020-00454-2.

50. Bensinger, S.J., Walsh, P.T., Zhang, J., Carroll, M., Parsons, R., Rathmell, J.C., Thompson, C.B., Burchill, M.A., Farrar, M.A., and Turka, L.A. (2004). Distinct IL-2 receptor signaling pattern in CD4+CD25+ regulatory T cells. J Immunol 172, 5287–5296. 10.4049/jimmunol.172.9.5287.

51. Schnell, A., Bod, L., Madi, A., and Kuchroo, V.K. (2020). The yin and yang of co-inhibitory receptors: toward anti-tumor immunity without autoimmunity. Cell Res 30, 285–299. 10.1038/s41422-020-0277-x.

52. Li, M.O., and Rudensky, A.Y. (2016). T cell receptor signalling in the control of regulatory T cell differentiation and function. Nature Reviews Immunology 16, 220–233. 10.1038/nri.2016.26.

53. Baine, I., Abe, B.T., and Macian, F. (2009). Regulation of T-cell tolerance by calcium/NFAT signaling. Immunol Rev 231, 225–240. 10.1111/j.1600-065X.2009.00817.x.

54. Martinez, G.J., Pereira, R.M., Aijo, T., Kim, E.Y., Marangoni, F., Pipkin, M.E., Togher, S., Heissmeyer, V., Zhang, Y.C., Crotty, S., et al. (2015). The Transcription Factor NFAT Promotes Exhaustion of Activated CDS T Cells. Immunity 42, 265–278. 10.1016/j.immuni.2015.01.006.

55. Kalekar, L.A., Schmiel, S.E., Nandiwada, S.L., Lam, W.Y., Barsness, L.O., Zhang, N., Stritesky, G.L., Malhotra, D., Pauken, K.E., Linehan, J.L., et al. (2016). CD4(+) T cell anergy prevents autoimmunity and generates regulatory T cell precursors. Nat Immunol 17, 304–314. 10.1038/ni.3331.

56. Van de Sande, B., Flerin, C., Davie, K., De Waegeneer, M., Hulselmans, G., Aibar, S., Seurinck, R., Saelens, W., Cannoodt, R., Rouchon, Q., et al. (2020). A scalable SCENIC workflow for single-cell gene regulatory network analysis. Nat Protoc 15, 2247–2276. 10.1038/s41596-020-0336-2.

57. Yamada, T., Park, C.S., Mamonkin, M., and Lacorazza, H.D. (2009). Transcription factor ELF4 controls the proliferation and homing of CDS+ T cells via the Kruppel-like factors KLF4 and KLF2. Nat Immunol 10, 618–626. 10.1038/ni.1730.

58. Wagle, M.V., Vervoort, S.J., Kelly, M.J., Van der Byl, W., Peters, T.J., Martin, B.P., Martelotto, L.G., Nüssing, S., Ramsbottom, K.M., Torpy, J.R., et al. (2021). Antigen-driven EGR2 expression is required for exhausted CDS T cell stability and maintenance. Nat Commun 12. ARTN 2782 10.1038/s41467-021-23044-9.

59. Trujillo-Ochoa, J.L., Kazemian, M., and Afzali, B. (2023). The role of transcription factors in shaping regulatory T cell identity. Nat Rev Immunol. 10.1038/s41577-023-00893-7.

60. Garg, G., Muschaweckh, A., Moreno, H., Vasanthakumar, A., Floess, S., Lepennetier, G., Oellinger, R., Zhan, Y.F., Regen, T., Hiltensperger, M., et al. (2019). Blimpl Prevents Methylation of and Loss of Regulatory T Cell Identity at Sites of Inflammation. Cell Reports 26, 1854-+. 10.1016/j.celrep.2019.01.070.

61. Wang, L., Liu, Y., Han, R., Beier, U.H., Thomas, R.M., Wells, A.D., and Hancock, W.W. (2013). Mbd2 promotes foxp3 demethylation and T-regulatory-cell function. Mol Cell Biol 33, 4106–4115. 10.1128/MCB.00144-13.

62. Kim, H.J., Barnitz, R.A., Kreslavsky, T., Brown, F.D., Moffett, H., Lemieux, M.E., Kaygusuz, Y., Meissner, T., Holderried, T.A., Chan, S., et al. (2015). Stable inhibitory activity of regulatory T cells requires the transcription factor Helios. Science 350, 334–339. 10.1126/science.aad0616.

63. Bending, D., and Zikherman, J. (2023). Nr4a nuclear receptors: markers and modulators of antigen receptor signaling. Curr Opin Immunol 81. ARTN 102285 10.1016/j.coi.2023.102285.

64. Rudensky, A.Y., Gavin, M., and Zheng, Y. (2006). FOXP3 and NFAT: partners in tolerance. Cell 126, 253–256. 10.1016/j.cell.2006.07.005.

65. Mucida, D., Pino-Lagos, K., Kim, G., Nowak, E., Benson, M.J., Kronenberg, M., Noelle, R.J., and Cheroutre, H. (2009). Retinoic acid can directly promote TGF-beta-mediated Foxp3(+) Treg cell conversion of naive T cells. Immunity 30, 471–472; author reply 472-473. 10.1016/j.immuni.2009.03.008.

66. Cunningham, T.J., and Duester, G. (2015). Mechanisms of retinoic acid signalling and its roles in organ and limb development. Nature Reviews Molecular Cell Biology 16, 110–123. 10.1038/nrm3932.

67. Casaletto, K.B., Elahi, F.M., Bettcher, B.M., Neuhaus, J., Bendlin, B.B., Asthana, S., Johnson, S.C., Yaffe, K., Carlsson, C., Blennow, K., et al. (2017). Neurogranin, a synaptic protein, is associated with memory independent of Alzheimer biomarkers. Neurology 89, 1782–1788. 10.1212/WNL.0000000000004569.

68. La Manno, G., Soldatov, R., Zeisel, A., Braun, E., Hochgerner, H., Petukhov, V., Lidschreiber, K., Kastriti, M.E., Lonnerberg, P., Furlan, A., et al. (2018). RNA velocity of single cells. Nature 560, 494–498. 10.1038/s41586-018-0414-6.

69. Bergen, V., Lange, M., Peidli, S., Wolf, F.A., and Theis, F.J. (2020). Generalizing RNA velocity to transient cell states through dynamical modeling. Nat Biotechnol 38, 1408–1414. 10.1038/s41587-020-0591-3.

70. Qiu, X., Hill, A., Packer, J., Lin, D., Ma, Y.A., and Trapnell, C. (2017). Single-cell mRNA quantification and differential analysis with Census. Nat Methods 14, 309–315. 10.1038/nmeth.4150.

71. Cousins, S.W., McCabe, M.M., Danielpour, D., and Streilein, J.W. (1991). Identification of transforming growth factor-beta as an immunosuppressive factor in aqueous humor. Invest Ophthalmol Vis Sci 32, 2201–2211.

72. Coombes, J.L., Siddiqui, K.R., Arancibia-Carcamo, C.V., Hall, J., Sun, C.M., Belkaid, Y., and Powrie, F. (2007). A functionally specialized population of mucosal CD103+ DCs induces Foxp3+ regulatory T cells via a TGF-beta and retinoic acid-dependent mechanism. J Exp Med 204, 1757–1764. 10.1084/jem.20070590.

73. Hong, S.W., Krueger, P.D., Osum, K.C., Dileepan, T., Herman, A., Mueller, D.L., and Jenkins, M.K. (2022). Immune tolerance of food is mediated by layers of CD4(+) T cell dysfunction. Nature 607, 762–768. 10.1038/s41586-022-04916-6.

74. Hoffman, L., Chandrasekar, A., Wang, X., Putkey, J.A., and Waxham, M.N. (2014). Neurogranin alters the structure and calcium binding properties of calmodulin. J Biol Chem 289, 14644–14655. 10.1074/jbc.Mll4.560656.

75. Iniguez, M.A., Morte, B., Rodriguez-Pena, A., Munoz, A., Gerendasy, D., Sutcliffe, J.G., and Bernal, J. (1994). Characterization of the promoter region and flanking sequences of the neuron-specific gene RC3 (neurogranin). Brain Res Mol Brain Res 27, 205–214. 10.1016/0169-328x(94)90002-7.

76. Dutta, D., Barr, V.A., Akpan, I., Mittelstadt, P.R., Singha, L.I., Samelson, L.E., and Ashwell, J.D. (2017). Recruitment of calcineurin to the TCR positively regulates T cell activation. Nat Immunol 18, 196–204. 10.1038/ni.3640.

77. McLane, L.M., Abdel-Hakeem, M.S., and Wherry, E.J. (2019). CDS T Cell Exhaustion During Chronic Viral Infection and Cancer. Annu Rev Immunol 37, 457–495. 10.1146/annurev-immunol-041015-055318.

78. Heng, J.S., Hackett, S.F., Stein-O’Brien, G.L., Winer, B.L., Williams, J., Goff, L.A., and Nathans, J. (2019). Comprehensive analysis of a mouse model of spontaneous uveoretinitis using single-cell RNA sequencing. P Natl Acad Sci USA 116, 26734–26744. 10.1073/pnas.191557lll6.

79. Lipski, D.A., Dewispelaere, R., Foucart, V., Caspers, L.E., Defrance, M., Bruyns, C., and Willermain, F. (2017). MHC class II expression and potential antigen-presenting cells in the retina during experimental autoimmune uveitis. J Neuroinflamm 14. 10.1186/sl2974-017-0915-5.

80. Okunuki, Y., Mukai, R., Nakao, T., Tabor, S.J., Butovsky, O., Dana, R., Ksander, B.R., and Connor, K.M. (2019). Retinal microglia initiate neuroinflammation in ocular autoimmunity. P Natl Acad Sci USA 116, 9989–9998. 10.1073/pnas.1820387116.

81. Nosko, A., Kluger, M.A., Diefenhardt, P., Melderis, S., Wegscheid, C., Tiegs, G., Stahl, R.A., Panzer, U., and Steinmetz, O.M. (2017). T-Bet Enhances Regulatory T Cell Fitness and Directs Control of Thl Responses in Crescentic GN. J Am Soc Nephrol 28, 185–196. 10.1681/ASN.2015070820.

82. Levine, A.G., Mendoza, A., Hemmers, S., Moltedo, B., Niec, R.E., Schizas, M., Hoyos, B.E., Putintseva, E.V., Chaudhry, A., Dikiy, S., et al. (2017). Stability and function of regulatory T cells expressing the transcription factor T-bet. Nature 546, 421–425. 10.1038/nature22360.

83. Paust, H.J., Riedel, J.H., Krebs, C.F., Turner, J.E., Brix, S.R., Krohn, S., Velden, J., Wiech, T., Kaffke, A., Peters, A., et al. (2016). CXCR3+ Regulatory T Cells Control THl Responses in Crescentic GN. J Am Soc Nephrol 27, 1933–1942. 10.1681/ASN.2015020203.

84. Luger, D., Silver, P.B., Tang, J., Cua, D., Chen, Z., Iwakura, Y., Bowman, E.P., Sgambellone, N.M., Chan, C.C., and Caspi, R.R. (2008). Either a Thl7 or a Thl effector response can drive autoimmunity: conditions of disease induction affect dominant effector category. J Exp Med 205, 799–810. 10.1084/jem.20071258.

85. Horai, R., Zarate-Blades, C.R., Dillenburg-Pilla, P., Chen, J., Kielczewski, J.L., Silver, P.B., Jittayasothorn, Y., Chan, C.C., Yamane, H., Honda, K., and Caspi, R.R. (2015). Microbiota-Dependent Activation of an Autoreactive T Cell Receptor Provokes Autoimmunity in an Immunologically Privileged Site. Immunity 43, 343–353. 10.1016/j.immuni.2015.07.014.

86. Fontenot, J.D., Rasmussen, J.P., Gavin, M.A., and Rudensky, A.Y. (2005). A function for interleukin 2 in Foxp3-expressing regulatory T cells. Nat Immunol 6, 1142–1151. 10.1038/nil263.

