## Supplemental Figure 1 (Fig. S1) for "Ocular immune privilege in action: the living eye imposes unique regulatory and anergic gene signatures on uveitogenic T cells"

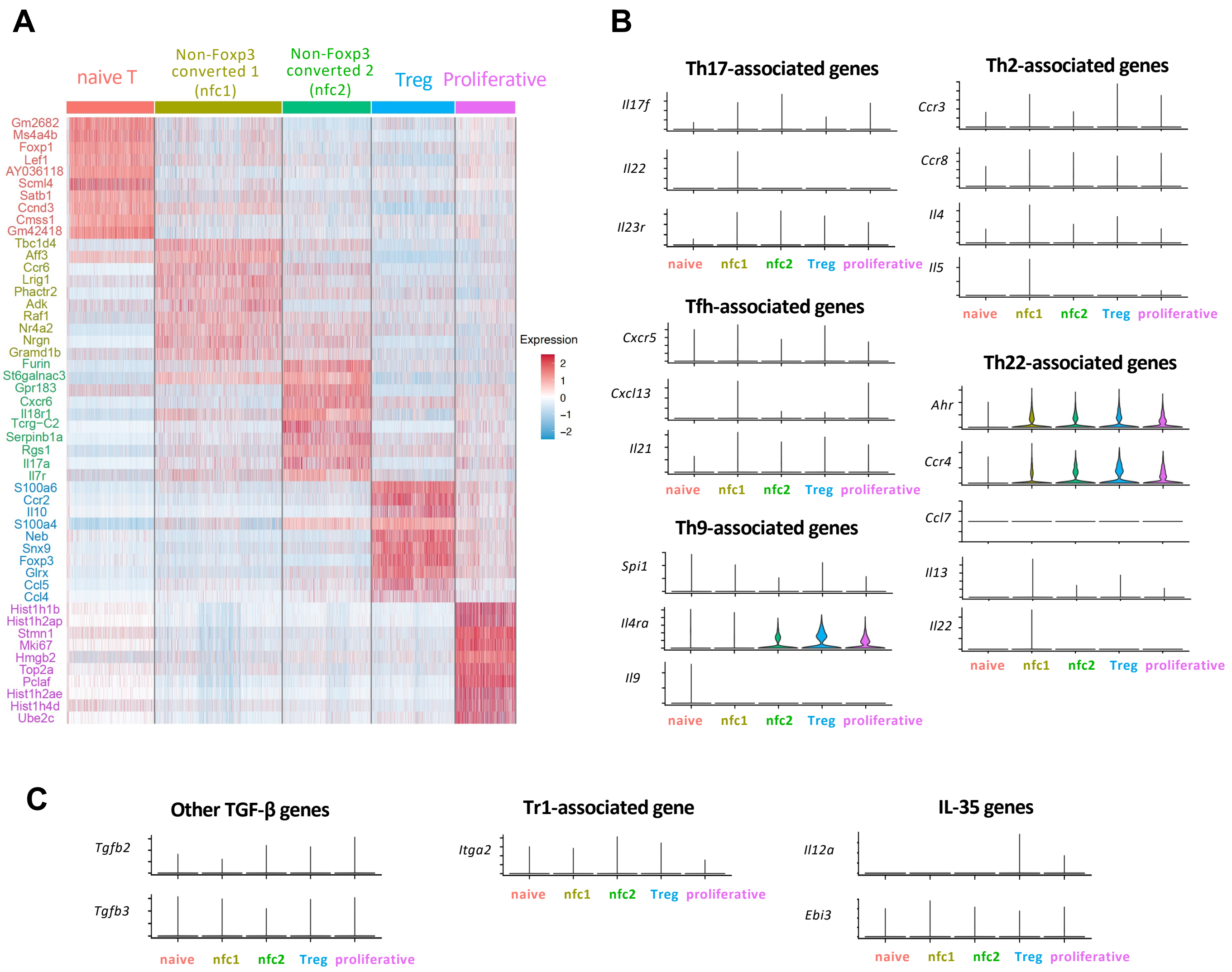


**Figure. S1: Transcriptional landscape suggests non-Foxp3-converted subsets are composed of Th-lineage-negative cells**

(A) Heatmap of the top 10 differentially expressed genes in each cluster *vs.* all other clusters, showing scaled expression of the top 10 genes in each cluster. (B) Violin plots showing expression of Th effector lineage-related transcription factors (TFs), cytokines, chemokines/chemokine receptors: Th17-related (*Csf2, Il17f, Il22*, and *Il23r)*, Th2-associated (*Ccr3, Ccr8, Il4, Il5, Il9),* Tfh-associated (*Cxcr5, Cxcl13, Il21),* Th9-associated (*Spi1*, encoding PU.1, *Il4ra*, *Il9),* and Th22-associated (*Ahr*, *Ccr4*, *Ccl7*, *Il13*, *Il22*). (C) Violin plots showing the absence of Tr1-associated gene *Itga2* (encoding CD49b), and genes for TGF-β2, TGF-β3 and IL-35 (*Il12a, Ebi3*).
